# Nrf2 regulates ICAM-1–mediated neutrophil extracellular trap formation after traumatic brain injury

**DOI:** 10.64898/2026.05.01.722360

**Authors:** P. M. Abdul Muneer, Saurav Bhowmick, Yemin A. Poovanthodi, Saleena Alikunju

## Abstract

Traumatic brain injury (TBI) induces persistent neurovascular inflammation that contributes to blood-brain barrier (BBB) disruption and secondary neurological dysfunction, but the mechanisms linking endothelial oxidative stress to leukocyte recruitment and neutrophil extracellular trap (NET) formation remain incompletely defined. Here, we investigated whether endothelial nuclear factor erythroid 2–related factor 2 (Nrf2) coordinates ICAM-1–dependent inflammatory responses after TBI. Using fluid percussion injury in wild-type, Nrf2-deficient, and ICAM-1-deficient mice, together with stretch-injured human brain microvascular endothelial cells, we examined oxidative stress, endothelial adhesion signaling, leukocyte recruitment, BBB integrity, NET formation, and neurological outcomes. TBI reduced endothelial Nrf2 activity and increased oxidative stress and ICAM-1 expression. Nrf2 deficiency further enhanced ICAM-1 expression, LFA-1/Mac-1–dependent leukocyte adhesion and transmigration, and BBB disruption, whereas Nrf2 activation attenuated these responses. ICAM-1 silencing or genetic deficiency rescued endothelial junctional disruption and reduced leukocyte recruitment, while recombinant ICAM-1 reproduced key features of barrier dysfunction. Nrf2 deficiency also increased neutrophil recruitment and NET formation, whereas ICAM-1 deficiency markedly suppressed these responses. Pharmacological and genetic inhibition of PAD4/NET formation reduced downstream endothelial injury. These findings identify an endothelial Nrf2–ICAM-1 axis linking oxidative stress to leukocyte recruitment and NET-mediated neurovascular injury after TBI.

## INTRODUCTION

Neuroinflammation following traumatic brain injury (TBI) involves activation of glial cells, release of pro-inflammatory cytokines and chemokines, upregulation of adhesion molecules, and recruitment of peripheral immune cells such as leukocytes (1). Studies from our laboratory and others have demonstrated that oxidative signaling is a central mechanism underlying TBI-associated disruption of the blood-brain barrier (BBB) and neurovascular impairments, which facilitate infiltration of inflammatory cells into the brain (2–7). Consequently, targeting oxidative stress represents a promising therapeutic strategy for mitigating TBI-associated secondary injury.

The transcription factor nuclear factor erythroid 2–related factor 2 (Nrf2), a member of the Cap ‘n’ Collar family, is a master regulator of cellular defense against oxidative stress through coordinated transcriptional activation of antioxidant, detoxification, and cytoprotective genes (8–13). In the central nervous system, Nrf2 confers robust neuroprotection by safeguarding neurons and astrocytes from toxic insults, regulating microglial activation, modulating inflammatory gene expression, and enhancing endogenous antioxidant enzyme activity (14–17). Beyond its canonical antioxidant role, emerging evidence highlights Nrf2 as a regulator of neurovascular inflammation through transcriptional control of endothelial adhesion molecules such as Vascular cell adhesion molecule-1 (VCAM-1) and intercellular adhesion molecule-1 (ICAM-1), thereby modulating leukocyte–endothelial interactions and vascular permeability (18–20). Therapeutic induction of Nrf2 therefore offers a mechanistically integrated and potentially safer alternative to exogenous antioxidant strategies by activating endogenous protective pathways with minimal off-target toxicity (21, 22).

Intercellular adhesion molecule-1 (ICAM-1) is a key endothelial adhesion protein that mediates leukocyte adhesion and transmigration during neuroinflammatory responses. Under basal conditions, ICAM-1 expression is low but is strongly upregulated by inflammatory stimuli such as tumor necrosis factor-α (TNF-α) (23, 24). ICAM-1 interacts with leukocyte integrins, including lymphocyte function-associated antigen 1 (LFA-1) and macrophage-1 antigen (Mac-1), promoting leukocyte adhesion, transmigration, and downstream inflammatory signaling cascades (25). Following TBI, infiltration of polymorphonuclear leukocytes (PMNs) contributes to neurovascular injury by activating endothelial adhesion molecules and stimulating the release of pro-inflammatory cytokines (26). Although Nrf2 has been implicated in the regulation of endothelial adhesion molecule expression and vascular inflammatory responses (1, 27, 28), whether loss of endothelial Nrf2 activity after TBI promotes ICAM-1–dependent leukocyte adhesion and transmigration, and whether this pathway contributes to subsequent neurovascular injury, remains unclear. Defining this relationship is particularly important because persistent endothelial-leukocyte interactions may provide a mechanistic link between post-traumatic oxidative stress and downstream inflammatory processes within the neurovascular unit.

Activated neutrophils release nuclear and granular contents to form neutrophil extracellular traps (NETs), web-like DNA structures implicated in TBI, stroke, and other neuroinflammatory conditions (29–33) NETs have also been associated with autoimmune disorders, cardiovascular and pulmonary diseases, inflammation, thrombosis, and cancer, underscoring their broad pathological relevance (34–37). ICAM-1–mediated leukocyte adhesion and transmigration may facilitate NET formation within the neurovascular unit, further amplifying inflammatory and vascular damage after brain injury (38). Although Nrf2 has been shown to regulate adhesion molecule expression and attenuate inflammation, its role in modulating ICAM- 1–dependent leukocyte transmigration and NET formation following TBI remains unclear.

Here, we tested the hypothesis that impaired endothelial Nrf2 activity after TBI establishes a redox-sensitive inflammatory state that activates ICAM-1-dependent leukocyte recruitment and promotes subsequent PAD4-dependent NET formation, thereby amplifying BBB disruption and secondary neurovascular injury. Using *in vivo* fluid percussion injury in wild-type, *Nrf2^−^/^−^* and *ICAM^−^/^−^* mice, and *in vitro* stretch injury in human brain microvascular endothelial cells, we manipulated Nrf2, ICAM-1, and peptidyl arginine deiminase-4 (PAD4) through pharmacological and genetic approaches. Our findings demonstrate that Nrf2 loss exacerbates ICAM-1-associated inflammatory responses, leukocyte recruitment, and NET formation following TBI, contributing to neurovascular injury and impaired recovery.

## RESULTS

### TBI suppresses Nrf2 signaling in cerebral microvascular endothelial cells and brain microvessels

To determine whether Nrf2 contributes to neurovascular protection after TBI, we first examined the expression and localization of total and phosphorylated Nrf2 (p-Nrf2) in brain microvessels isolated from the pericontusional cortex. Double immunofluorescence staining revealed that both Nrf2 and p-Nrf2 co-localized with the endothelial marker von Willebrand factor (vWF).

While total Nrf2 was predominantly cytoplasmic, p-Nrf2 localized mainly to the nucleus, consistent with activation-dependent nuclear translocation. Following injury, however, both total and phosphorylated Nrf2 signals were markedly reduced (∼ 50%; *p<0.001*) in endothelial cells (Fig. 1A-C). These observations were confirmed biochemically. Western blotting demonstrated a significant reduction (more than 50%; *p<0.001*) in Nrf2 and p-Nrf2 protein levels in the pericontusional cortex in injured wild-type (WT) mice compared with uninjured controls, accompanied by a parallel decrease in Nrf2 mRNA expression (∼37%; *p<0.001*) as assessed by qRT-PCR (Fig. 1D-F). As expected, Nrf2 and p-Nrf2 were undetectable in *Nrf2^−^/^−^* mice.

**Figure 1:**
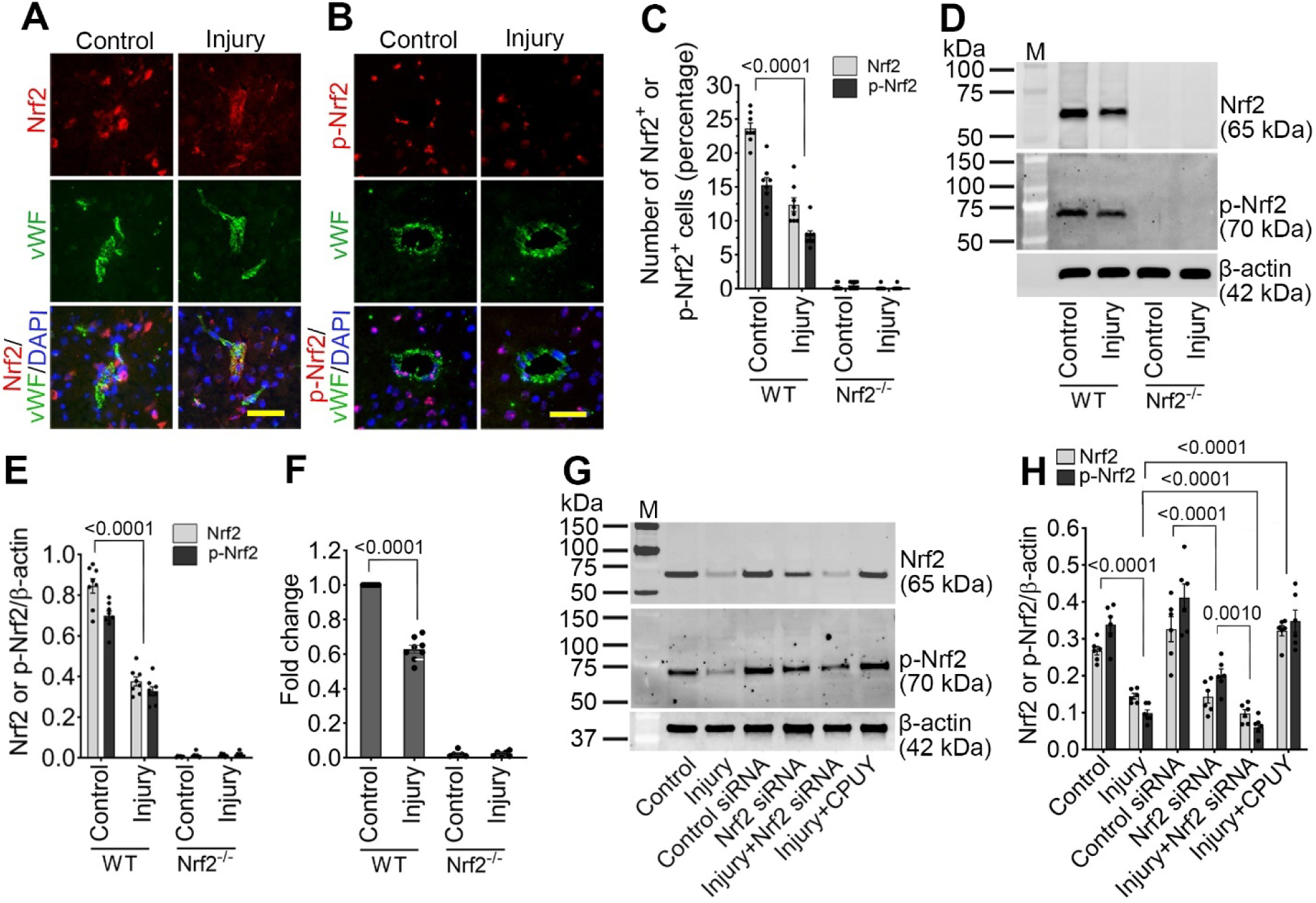
TBI suppresses endothelial Nrf2 signaling *in vivo* and *in vitro*. (**A-B**) Representative immunofluorescence images showing Nrf2 and phosphorylated Nrf2 (p-Nrf2) (red) colocalized with the endothelial marker von Willebrand factor (vWF; green) and nuclei counterstained with DAPI (blue) in the pericontusional cortex in WT control and injured mice 24 h following 25 psi fluid percussion injury (FPI). Scale bar = 50 µm. (**C**) Bar graph showing the percentage of Nrf2 and p-Nrf2 positive cells (n = 6/group). (**D**) Western blot analysis of Nrf2, p-Nrf2, and β-actin in cortical tissue lysates from WT and *Nrf2^−^/^−^* control and injured mice 24 h after 25 psi FPI. (**E**) Bar graph showing quantification of Nrf2 and p-Nrf2 protein levels normalized to β-actin (n = 6/group). (**F**) mRNA expression of Nrf2 determined by RT-qPCR in WT and *Nrf2^−^/^−^* mice 24 h after 25 psi FPI. Bar graph represents fold change relative to WT control (n = 6/group). (**G**) Western blot analysis of Nrf2, p-Nrf2, and β-actin in human brain microvascular endothelial cells (HBMECs) 24 h after 3.0 psi stretch injury following treatment with control siRNA, Nrf2 siRNA, or CPUY192018. (**H**) Bar graph shows quantification of Nrf2 and p-Nrf2 protein levels normalized to β-actin (n = 4/group). All values are expressed as mean ± SEM. Statistical analysis was carried out by two-way ANOVA (for **C**, **E**, and **F**) and one-way ANOVA (for **H**) with Dunnett’s post hoc test. p < 0.05 statistically significant. ‘ns’ is not significant.

To assess whether mechanical injury directly impairs endothelial Nrf2 signaling, human brain microvascular endothelial cells (hBMVECs) were subjected to stretch injury following pretreatment with control siRNA, Nrf2 siRNA, or the Keap1-Nrf2 interaction inhibitor CPUY192018. Stretch injury resulted in a significant reduction in Nrf2 (∼37%; *p<0.001*) and p- Nrf2 (∼75%; *p<0.001*) protein expression at 24 h post-injury. Nrf2 knockdown further exacerbated this reduction in stretch-injured samples, whereas pharmacological disruption of Keap1–Nrf2 binding significantly increased both total and phosphorylated Nrf2 levels (∼1.9 and 4.3-fold respectively; *p<0.001*) compared with untreated injured cells (Fig. 1G and H). These data demonstrate that TBI suppresses endothelial Nrf2 signaling both *in vivo* and *in vitro*, and that this impairment can be reversed by pharmacological Nrf2 activation.

### Impaired Nrf2 signaling suppresses antioxidant gene expression and enhances oxidative stress following injury

Because Nrf2 mediates cellular antioxidant defense through activation of antioxidant response element (ARE)-driven genes (39), we next examined the impact of injury-induced Nrf2 suppression on key antioxidant enzymes. In stretch-injured hBMVECs, protein expression of GPx1, GSTm1, HO-1, and NQO1 was reduced to approximately one-third: GPx1 and NQO1, one-fourth: HO-1, and one-half: GSTm1 of uninjured levels. As expected, Nrf2 knockdown suppressed the expression of all four proteins in control and stretch-injured samples, whereas CPUY192018 treatment restored their expression to near control levels (Fig. 2A-C). Consistent with these findings, ARE-driven transcriptional activity assessed using an NQO1-ARE luciferase reporter was reduced by ∼60% (*p<0.001*) in injured hBMVECs. Nrf2 silencing further diminished ARE activity in both injured and uninjured cells, while CPUY192018 significantly enhanced (∼3.2-fold; *p<0.001*) ARE activity in injured cells (Fig. 2D).

**Figure 2:**
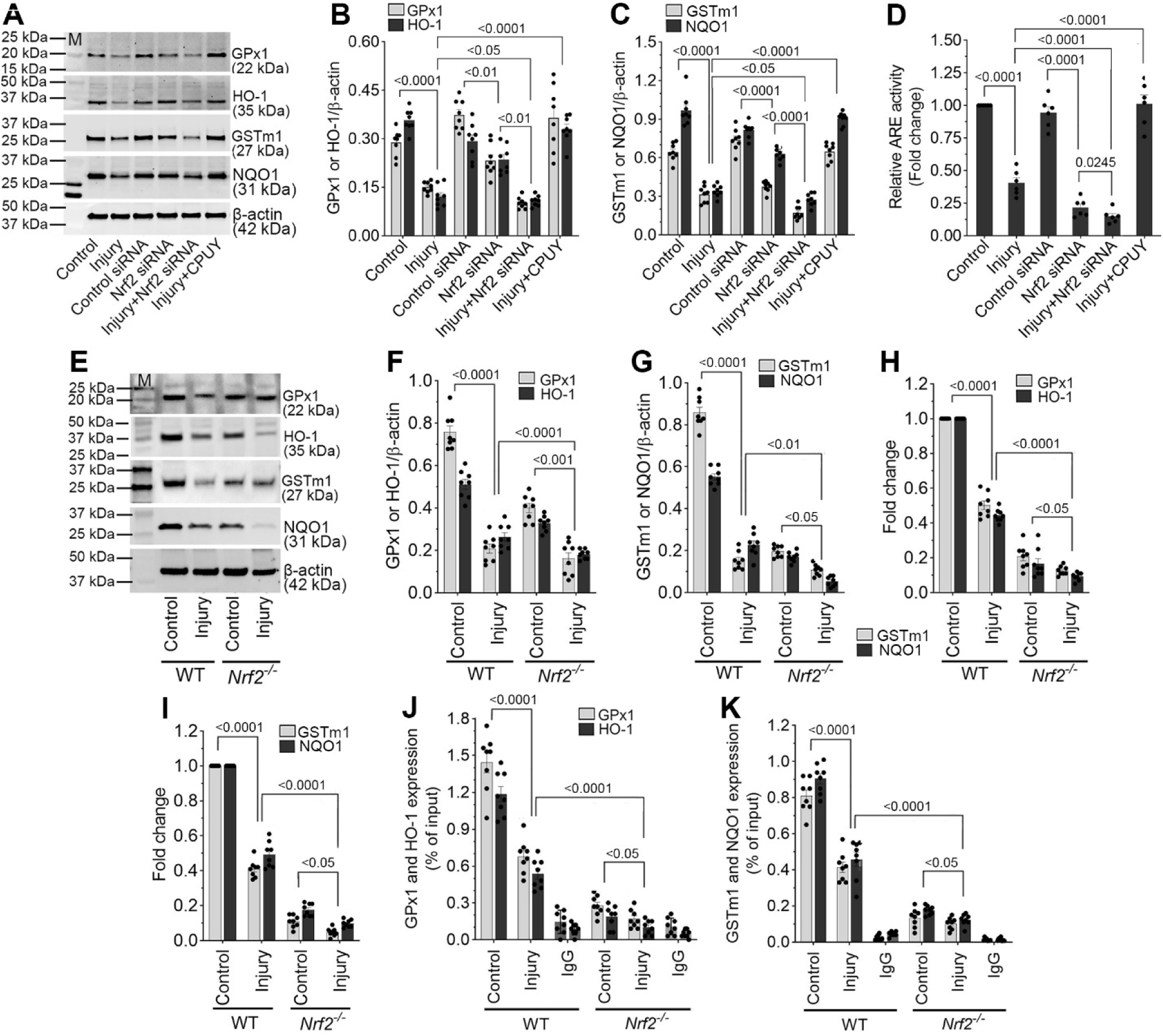
Impaired Nrf2 suppresses antioxidant gene expression following injury. (**A**) Western blot analysis of GPx1, HO-1, GSTm1, NQO1, and β-actin in human brain microvascular endothelial cells (hBMVECs) 24 h after 3.0 psi stretch injury following treatment with control siRNA, Nrf2 siRNA, or CPUY192018. (**B, C**) Bar graphs showing quantification of GPx1, HO-1 (B), GSTm1, and NQO1 (C) protein levels normalized to β-actin (n = 8/group). (**D**) ARE-driven transcriptional activity measured using an NQO1-ARE luciferase reporter assay in control and injured hBMVECs with Nrf2 siRNA or CPUY192018 treatment (n = 6/group). (**E**) Western blot analysis of GPx1, HO-1, GSTm1, NQO1, and β-actin in pericontusional cortical tissue from WT and *Nrf2^−^/^−^* mice 24 h after 25 psi fluid percussion injury (FPI). (**F, G**) Bar graphs showing quantification of antioxidant protein expression normalized to β-actin (n = 8/group). (**H, I**) RT-qPCR analysis of GPx1, HO-1 (H), GSTm1, and NQO1 (I) mRNA levels in cortical tissue from WT and *Nrf2^−^/^−^* mice 24 h after FPI (n = 8/group). Data are presented as fold change relative to WT control. (**J, K**) Chromatin immunoprecipitation (ChIP)-qPCR analysis showing Nrf2 binding to ARE regions within the promoters of GPx1, HO-1 (J), GSTm1, and NQO1 (K) in WT and *Nrf2^−^/^−^* mice following injury (n = 8/group). Quantification of Nrf2 occupancy at ARE sites demonstrating significant reduction after injury in WT mice and negligible binding in *Nrf2^−^/^−^* mice. All values are expressed as mean ± SEM. Statistical analysis was performed using one-way in **B-D** or two-way ANOVA in **F-K** followed by Dunnett’s post hoc test. p < 0.05 statistically significant.

*In vivo* analyses yielded similar results. Western blotting of pericontusional cortical tissue revealed marked reductions in GPx1 (∼one-third), GSTm1 (∼one-eighth), HO-1 (one-half), and NQO1 (one-fourth) expression following injury in WT mice. As expected, these antioxidant proteins were minimally expressed in *Nrf2^−^/^−^* mice in control and injured mice compared to their representative WT groups (Fig. 2E-G). Correspondingly, qRT-PCR analyses demonstrated significant reductions (∼one-half) in antioxidant gene transcripts after injury in WT mice, with consistently low basal expression in *Nrf2^−^/^−^* mice (Fig. 2H and I).

Chromatin immunoprecipitation-qPCR (ChIP-qPCR) further confirmed direct binding of Nrf2 to ARE regions within the promoters of GPx1, GSTm1, HO-1, and NQO1 in WT mice. Injury significantly reduced (*p<0.001*) Nrf2 occupancy at these sites, whereas binding was negligible in *Nrf2^−^/^−^* mice (Fig. 2J and K). Together, these data establish that TBI-induced impairment of Nrf2 signaling directly suppresses antioxidant gene transcription.

Given the reduction in antioxidant defenses, we next examined markers of oxidative and nitrosative stress. In stretch-injured hBMVECs, expression of NADPH oxidase-1 (NOX1, 1.6- fold) and inducible nitric oxide synthase (iNOS, 2.3-fold), along with their downstream oxidative products 4-hydroxynonenal (4-HNE, 5.4-fold) and 3-nitrotyrosine (3-NT, 7.4-fold), was significantly increased. Nrf2 knockdown further augmented these responses, whereas CPUY192018 treatment markedly reduced expression of all four markers (Fig. 3A-C).

**Figure 3:**
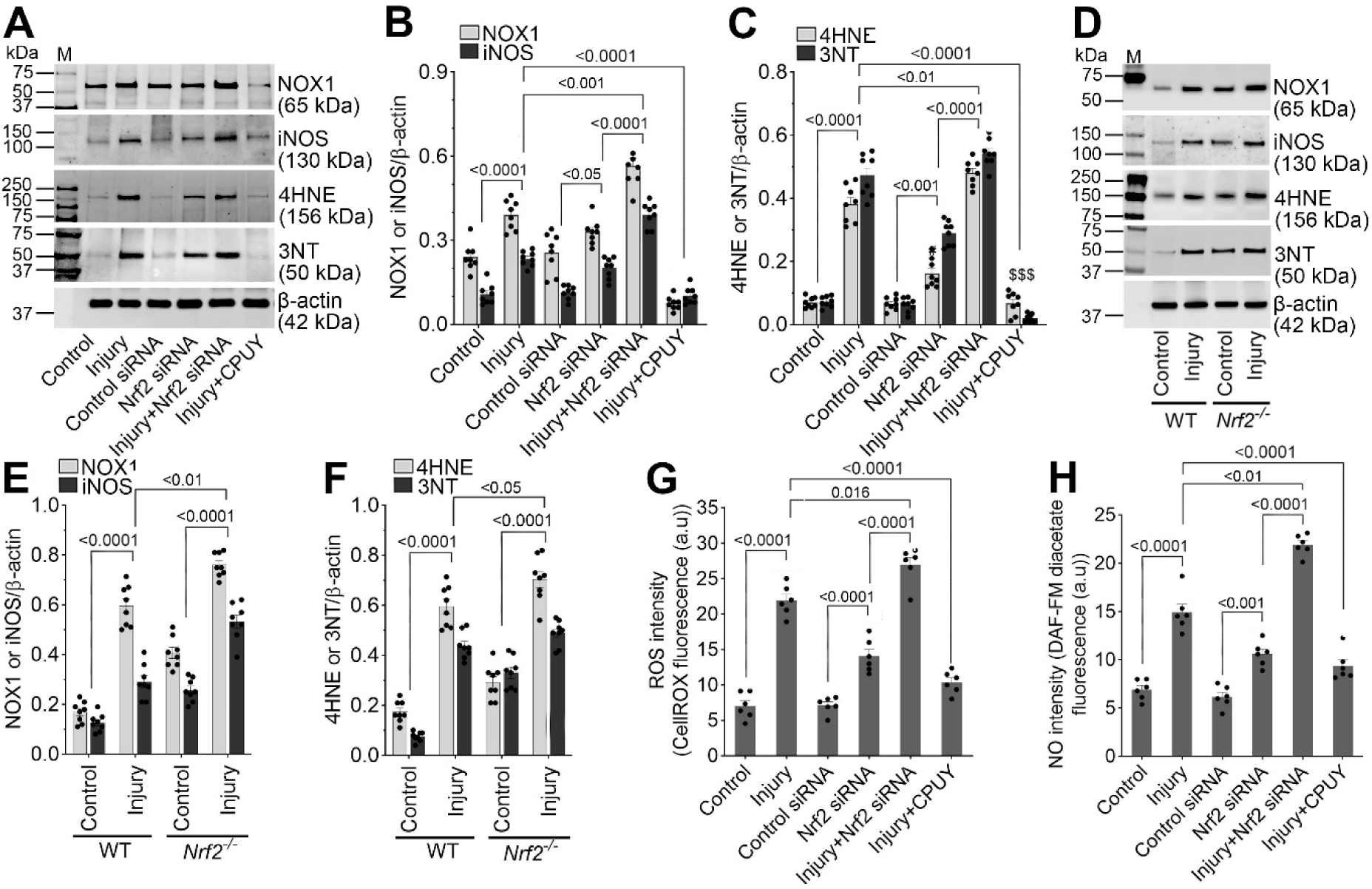
Nrf2 loss increases oxidative stress in stretch-injured hBMVECs and injured mice brain. (**A**) Western blot analysis of NOX1, iNOS, 4-HNE, 3-NT, and β-actin in human brain microvascular endothelial cells (hBMVECs) 24 h after 3.0 psi stretch injury following treatment with control siRNA, Nrf2 siRNA, or CPUY192018. (**B, C**) Bar graphs showing quantification of NOX1 and iNOS (B), and 4-HNE and 3-NT (C) protein levels normalized to β-actin (n = 8/group). (**D**) Western blot analysis of NOX1, iNOS, 4-HNE, 3-NT, and β-actin in pericontusional cortical tissue from WT and *Nrf2^−^/^−^* mice 24 h after 25 psi fluid percussion injury (FPI). (**E, F**) Bar graphs showing quantification of NOX1 and iNOS (E), and 4-HNE and 3-NT (F) protein levels normalized to β-actin (n = 8/group). (**G**) Representative images and quantification of intracellular reactive oxygen species (ROS) levels measured using CellROX Green fluorescence in control and stretch-injured hBMVECs with control siRNA, Nrf2 siRNA or CPUY192018 treatment (n = 6/group). (**H**) Quantification of nitric oxide (NO) production assessed by DAF-FM diacetate fluorescence in control and stretch-injured hBMVECs with control siRNA, Nrf2 siRNA or CPUY192018 treatment (n = 6/group). All values are expressed as mean ± SEM. Statistical analysis was performed using one-way ANOVA in **B, C, G** and **H**; and two-way ANOVA in **E** and **F**, followed by Dunnett’s post hoc test. *p < 0.05* statistically significant.

*In vivo*, injured WT mice exhibited elevated NOX1 (3.5-fold), iNOS (2.3-fold), 4-HNE (3.4-fold), and 3-NT (5.8-fold) levels in cortical tissue, which were further exacerbated in *Nrf2^−^/^−^* mice. Notably, uninjured *Nrf2^−^/^−^* mice displayed higher basal oxidative stress marker expression compared with WT controls (Fig. 3D-F).

Direct quantification of reactive oxygen species (ROS) using CellROX Green revealed a ∼3.0-fold increase in ROS levels following stretch injury, while nitric oxide production assessed using DAF-FM diacetate increased ∼2.0-fold. Nrf2 silencing further elevated ROS and NO levels, whereas CPUY192018 restored them to near baseline (Fig. 3G and H). These findings identify Nrf2 as a critical regulator of oxidative stress within the injured neurovascular unit.

### Nrf2 deficiency promotes ICAM-1 and leukocyte adhesion receptor activation

We next investigated whether Nrf2-mediated redox regulation influences endothelial activation. Immunohistochemical analysis revealed robust induction (3.3-fold) of intercellular adhesion molecule-1 (ICAM-1) in cortical microvessels following injury in WT mice. ICAM-1 expression was significantly greater in *Nrf2^−^/^−^* mice under both basal and injured conditions, whereas ICAM-1 was absent in *ICAM-1^−^/^−^* mice (Fig. 4A and B).

**Figure 4.**
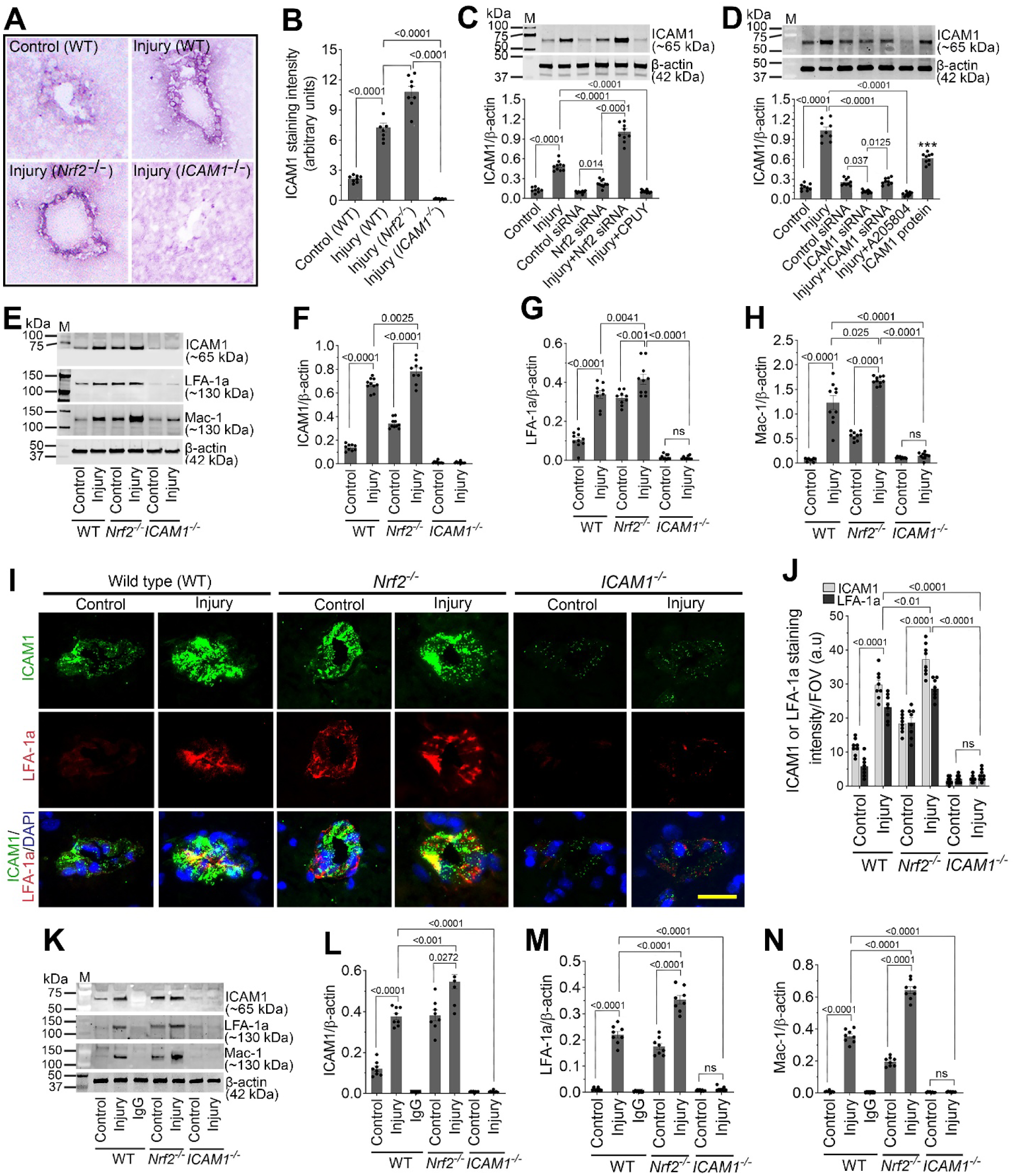
Nrf2 negatively regulates ICAM-1 and leukocyte adhesion receptors in hBMVECs and cortical microvessels. (**A-B**) Representative immunohistochemical images (A) and quantification (B) of ICAM-1 expression in cortical microvessels of WT, *Nrf2^−^/^−^*, and *ICAM-1^−^/^−^* mice 24 h after injury (n = 6/group). (**C**) Western blot analysis of ICAM-1 and β-actin in stretch-injured hBMVECs 24 h post-injury with control siRNA, Nrf2 siRNA, or CPUY192018 treatment (n = 10/group). (**D**) Western blot analysis of ICAM-1 and β-actin in stretch-injured hBMVECs 24 h post-injury with control siRNA, ICAM-1 siRNA, or A205804 (ICAM-1 inhibitor) treatment (n = 9/group). (**E**) Western blot expression level of ICAM-1, leukocyte adhesion receptors LFA-1 and Mac-1 and β-actin in pericontusional cortical tissue of WT, *Nrf2^−^/^−^* and *ICAM-1^−^/^−^* mice 24 h after injury. (**F-H**) Bar graphs showing quantification of ICAM-1 (F), leukocyte adhesion receptors LFA-1 (G) and Mac-1 (H) in pericontusional cortical tissue of WT, *Nrf2^−^/^−^* and *ICAM-1^−^/^−^* mice 24 h after injury (n = 9/group). (**I-J**) Double immunofluorescence showing co-localization of LFA-1 with ICAM-1 in cortical microvessels, particularly enhanced in *Nrf2^−^/^−^* mice (I) and corresponding quantification (J) (n = 8/group). (**K-N**) Immunoprecipitation demonstrating direct ICAM-1 interactions with LFA-1 (k, m) and Mac-1 (k, n) in cortical tissue (n = 8/group). All values are expressed as mean ± SEM. Statistical analysis was performed using one-way ANOVA in **B-D** or two-way ANOVA in **F-H, J, L-N** followed by Dunnett’s post hoc test*. p < 0.05* statistically significant. *\*\*\*P < 0.001* versus control in **D**. ‘ns’ is not significant.

In stretch-injured hBMVECs, ICAM-1 expression increased markedly (*p<0.001*) at 24 h post-injury. Nrf2 knockdown further enhanced ICAM-1 expression, while CPUY192018 significantly suppressed it (Fig. 4C). Direct silencing (with ICAM1 siRNA) or pharmacological inhibition (using A205804) of ICAM-1 significantly reduced its injury-induced expression, confirming specificity (Fig. 4D).

*In vivo*, ICAM-1 expression increased ∼4.8-fold in injured WT mice and was further elevated in injured *Nrf2^−^/^−^* mice. Notably, uninjured *Nrf2^−^/^−^* mice exhibited elevated basal ICAM- 1 levels compared with WT controls (Fig. 4E and F). Consistent with ICAM-1 activation, expression of its leukocyte receptors LFA-1 and Mac-1 was significantly increased (*p<0.001*) following injury in WT mice and further augmented in *Nrf2^−^/^−^* mice, while remaining minimal in *ICAM-1^−^/^−^* animals (Fig. 4G and H). Double immunofluorescence confirmed co-localization of LFA-1 with ICAM-1 in cortical microvessels, particularly in *Nrf2^−^/^−^* mice (Fig. 4I and J). Immunoprecipitation studies demonstrated direct ICAM-1 interactions with both LFA-1 and Mac- 1, which were enhanced by injury and further increased in *Nrf2^−^/^−^* mice (Fig. 4K-N).

### Nrf2 deficiency promotes vascular dysfunction and blood-brain barrier disruption after injury

Immunofluorescence and immunoblot analyses revealed significant loss of tight junction proteins occludin, claudin-5, and ZO-1 (approximately one-tenth, one-eight, and one-tenth, respectively) following stretch injury in hBMVECs. Nrf2 knockdown exacerbated this loss, whereas CPUY192018 restored tight junction protein expression. Importantly, ICAM-1 silencing, or inhibition prevented injury-induced tight junction disruption, while recombinant ICAM-1 reproduced the injury phenotype (Fig. 5A-E).

**Figure 5.**
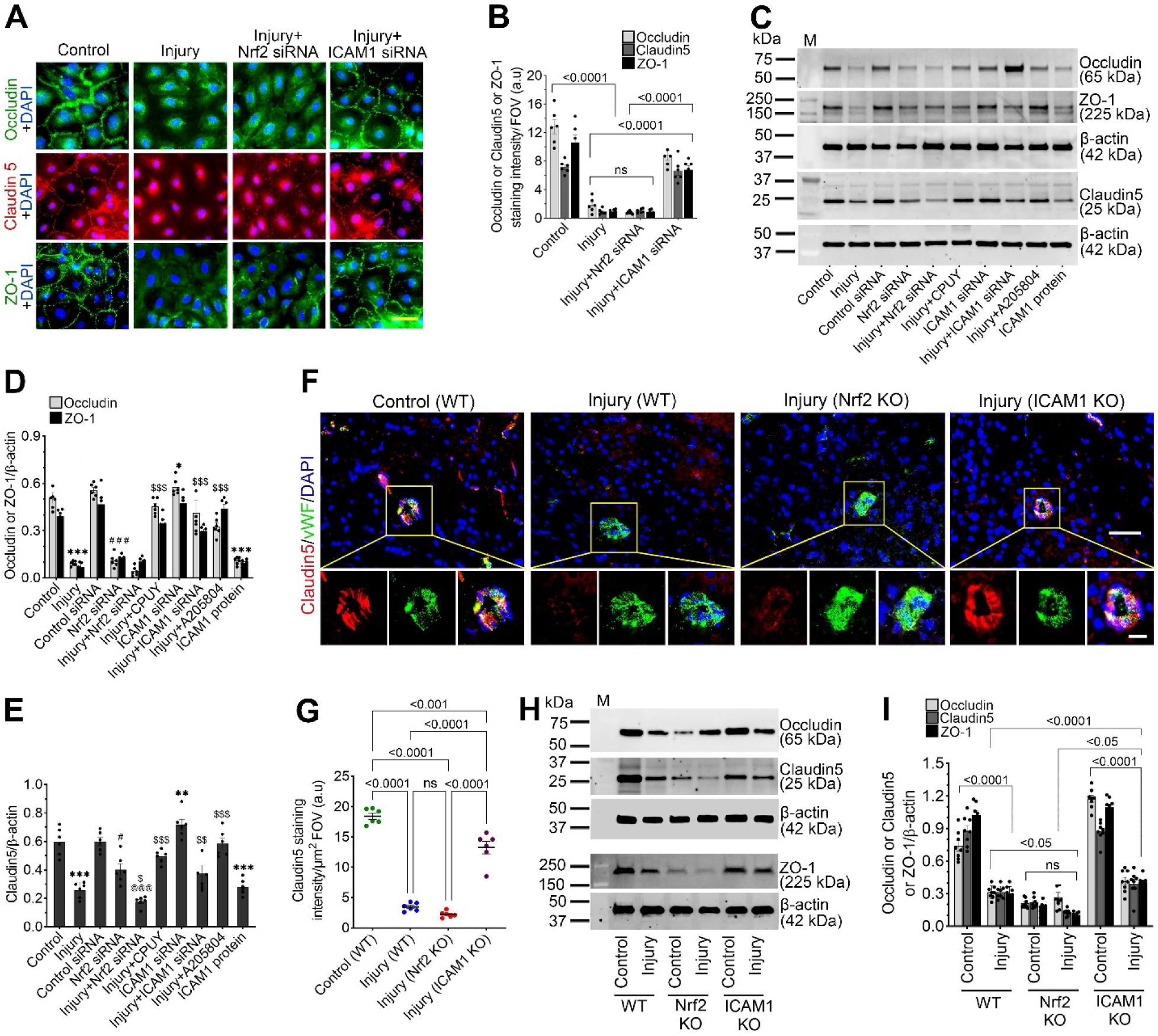
Nrf2 deficiency causes vascular dysfunction following traumatic brain injury. (**A-B**) Representative immunofluorescence staining (A) and quantification (B) of tight junction proteins Occludin (green), Claudin-5 (red), and ZO-1 (green) merged with DAPI (blue) in human brain microvascular endothelial cells (hBMVECs) following 3.0 psi stretch injury with or without Nrf2 or ICAM-1 siRNA treatment (n = 6/group). (**C-E**) Western blot analysis of Occludin, ZO-1, and Claudin-5 with β-actin as a loading control, in cell lysates collected 24 h after 3.0 psi stretch injury in hBMVECs treated with control siRNA, Nrf2 siRNA, the Nrf2 activator CPUY, ICAM-1 siRNA, the ICAM-1 inhibitor A2015804, or recombinant ICAM-1 protein. Bar graphs represent densitometric quantification of Occludin and ZO-1 (D) and Claudin-5 (E) normalized to β-actin (n = 6/group). (**F-G**) Representative immunofluorescence staining (F) and quantification (G) of Claudin-5 (red) merged with the endothelial marker vWF (green) and DAPI (blue) in cortical brain sections from uninjured and injured groups of WT, *Nrf2^−^/^−^* and *ICAM-1^−^/^−^* 24 hr after 25 psi fluid percussion injury (FPI) (n = 6/group). (**H-I**) Western blot analysis of Occludin, Claudin-5, and ZO-1, with β-actin as a loading control, in cortical brain tissue lysates from WT, *Nrf2^−^/^−^* and *ICAM-1^−^/^−^* mice following 15 psi FPI. Bar graphs (I) show densitometric quantification of Occludin, Claudin-5, and ZO-1 normalized to β-actin (n = 8/group). All values are expressed as mean ± SEM. Statistical analysis was performed using one-way ANOVA in **B**, **D**, **E**, and **G** or two-way ANOVA in **I** followed by Dunnett’s post hoc test. *p < 0.05* statistically significant. *\*P< 0.05, **P< 0.01, ***P < 0.001* versus control; *^#^P < 0.05, ^###^P < 0.001* versus control siRNA; *^@@@^P < 0.001* versus Nrf2 siRNA; *^$^P < 0.05, ^$$^P < 0.01, ^$$$^P < 0.001* versus injury in **D** and **E**. ‘ns’ is not significant.

*In vivo*, claudin-5 expression was reduced in injured WT mice and further decreased in *Nrf2^−^/^−^* mice, whereas *ICAM-1^−^/^−^* mice showed preservation of tight junction integrity (Fig. 5F-I). These findings suggest that Nrf2 deficiency is associated with increased ICAM-1 expression and contributes to tight junction disruption following TBI.

Beyond tight junctions, injury significantly reduced expression of the adherent junction protein N-cadherin and the gap junction protein connexin-43 in hBMVECs. Nrf2 knockdown exacerbated this reduction, while CPUY192018 restored expression. ICAM-1 silencing prevented injury-induced loss of both proteins, whereas recombinant ICAM-1 mimicked the injury response (Fig. 6A-D). *In vivo* analyses confirmed marked reductions in N-cadherin and connexin-43 following injury in WT mice, with further suppression in *Nrf2^−^/^−^* mice and preservation in *ICAM-1^−^/^−^* animals (Fig. 6E-G).

**Figure 6.**
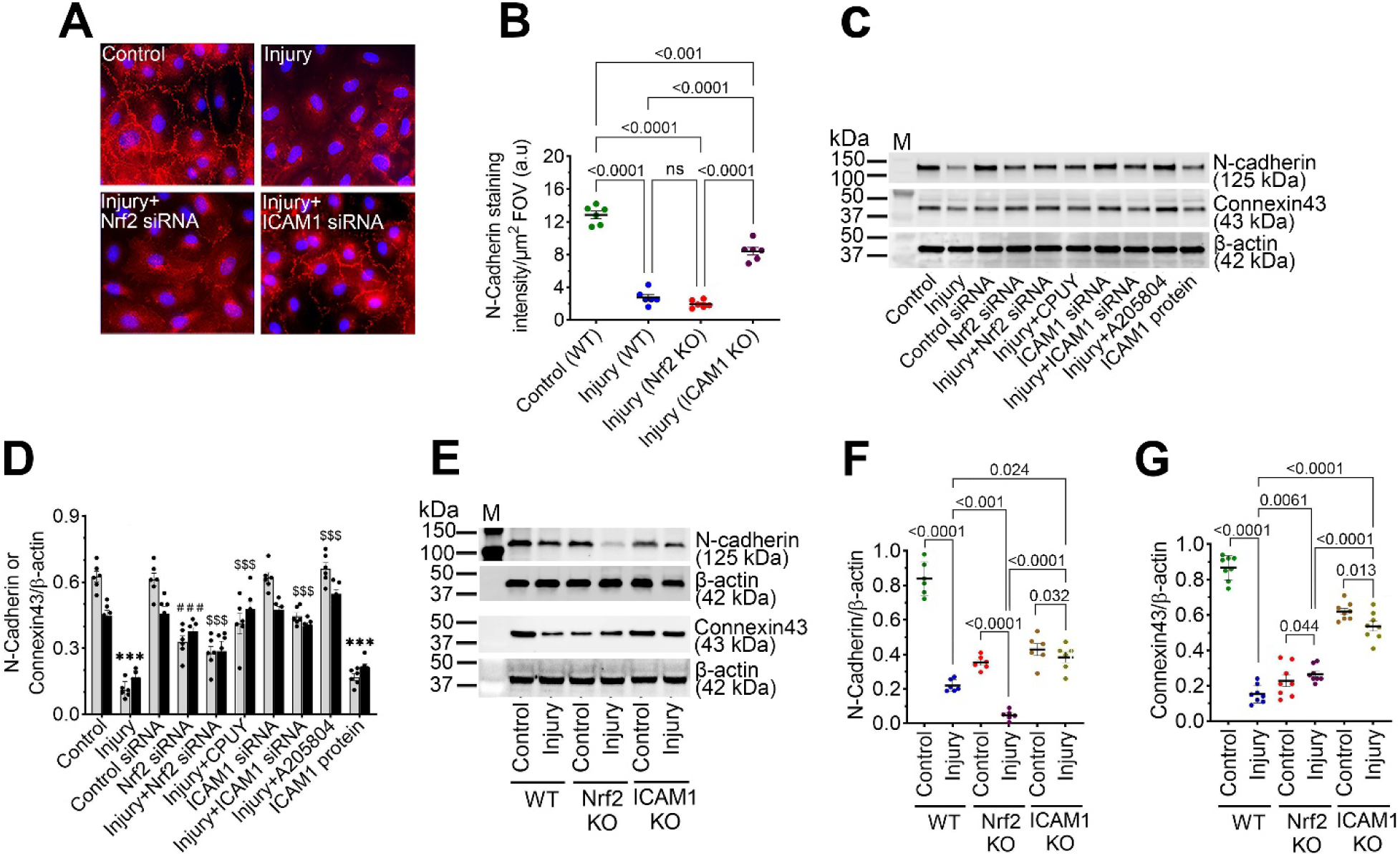
Nrf2 and ICAM-1 regulate junctional protein expression *in vitro* and *in vivo*. (**A-B**) Representative immunofluorescence staining (A) and quantification (B) of N-cadherin (red) with nuclear counterstain DAPI (blue) in hBMVEC following 3.0 psi stretch injury with or without Nrf2 or ICAM-1 siRNA treatment (n = 6/group). Scale bar = 40 µm. (**C-D**) Western blot analysis of N-cadherin, Connexin-43, and β-actin in lysates from stretch-injured hBMVEC cultures 24 h after treatment with control siRNA, Nrf2 siRNA, the Nrf2 activator CPUY, ICAM-1 siRNA, the ICAM-1 inhibitor A2015804, or recombinant ICAM-1 protein. Bar graphs represent densitometric quantification of N-cadherin (**D**) normalized to β-actin (n = 6/group). (**E-G**) Western blot analysis of N-cadherin and Connexin-43 with β-actin as a loading control in cortical brain tissue lysates from WT, *Nrf2^−^/^−^* and *ICAM-1^−^/^−^* mice following 15 psi FPI. Bar graphs show densitometric quantification of N-cadherin (**F**) and connexin-43 (**G**) normalized to β-actin (n = 6/group). All values are expressed as mean ± SEM. Statistical analysis was performed using one-way ANOVA in **B** and **D,** and two-way ANOVA in **F** and **G** followed by Dunnett’s post hoc test. Statistical analysis was performed using followed by Dunnett’s post hoc test. *p < 0.05* statistically significant. *\*\*\*P < 0.001* versus control; *^###^P < 0.001* versus control siRNA; *^$$$^P < 0.001* versus injury in **D**. ‘ns’ is not significant.

Functional assessment using an *in vitro* BBB model demonstrated significantly increased FITC-dextran permeability following stretch injury, which was further enhanced by Nrf2 knockdown. CPUY192018 or ICAM-1 inhibition significantly reduced permeability, whereas recombinant ICAM-1 increased leakiness (Supplementary Fig. 1A). *In vivo* tracer studies using sodium fluorescein and Evans blue revealed increased BBB permeability following injury in WT mice, which was exacerbated in *Nrf2^−^/^−^* mice but attenuated in *ICAM-1^−^/^−^* animals (Supplementary Fig. 1B and C).

### Nrf2 regulates ICAM-1–dependent leukocyte adhesion, recruitment, and transmigration

*In vitro* transmigration assays showed increased monocyte adhesion and migration (1.66-fold) across injured hBMVEC monolayers, which was further enhanced by Nrf2 knockdown and significantly reduced by CPUY192018 or ICAM-1 inhibition (Fig. 7A). GFP-monocyte assays confirmed increased adhesion at disrupted endothelial junctions in Nrf2-deficient conditions, while ICAM-1 silencing prevented monocyte adhesion (Fig. 7B and C).

**Figure 7.**
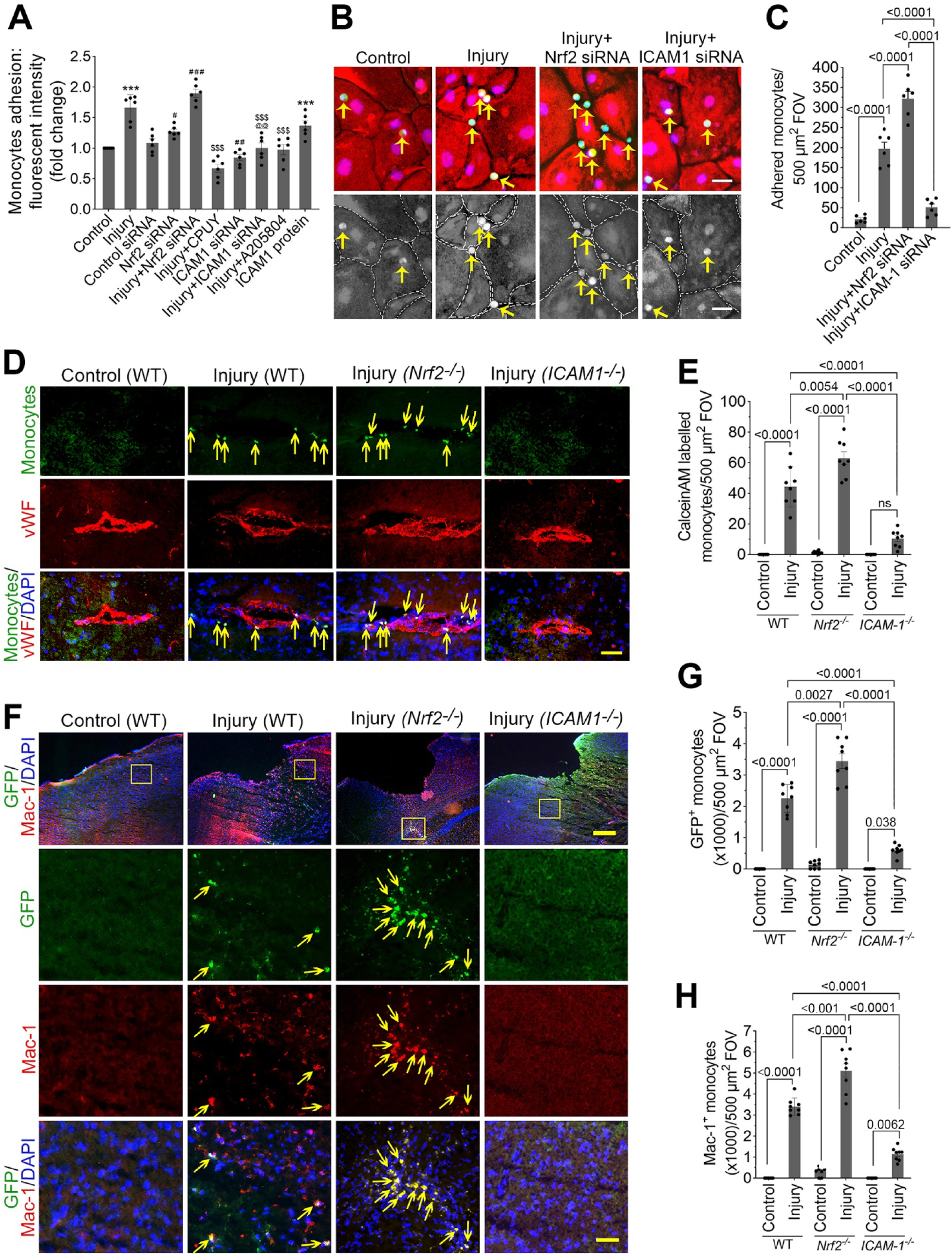
Nrf2 regulation of ICAM-1-mediated monocyte/macrophage adhesion and transmigration across the injured blood-brain barrier. **(A)** *In vitro* transmigration assay assessing monocyte adhesion and migration across injured human brain microvascular endothelial cell (hBMVEC) monolayers following Nrf2 knockdown, treatment with the Nrf2 activator CPUY192018, or ICAM-1 inhibition (n = 6/group). **(B-C)** Representative images and quantification of GFP-labeled monocyte adhesion (green) to injured hBMVEC monolayers (red). The nuclei were counter stained with DAPI (blue). Monocyte adhesion at endothelial junctions was evaluated under Nrf2-deficient conditions and after ICAM-1 silencing. Arrows show the adhered monocytes on the hBMVEC monolayer (n = 6/group). Scale bar = 10 µm. **(D-E)** Adhesion and transmigration of Calcein-AM–labeled macrophages in the perivascular space of wild-type (WT), *Nrf2^−^/^−^* and *ICAM-1^−^/^−^* mice following injury. Bar graph (E) shows the quantification of adhered and transmigrated macrophages in the perivascular space (n = 8/group). Scale bar = 100 µm. (**F-H**) Transmigration of leukocyte analysis by infusing cultured GFP monocytes (green) through the common carotid artery of WT, *Nrf2^−^/^−^* and *ICAM-1^−^/^−^* mice 24 h after injury and co-localized with immunostaining images of Mac-1 (red) and DAPI (blue). Panels in second through fourth rows are selected and enlarged areas of first row. Scale bar: 400 µm in the panels of first row and 80 µm in the panels of second through fourth rows. (**G-H**) Quantitative analysis of the number of infused GFP (G) and Mac-1 (H) positive cells. All values are expressed as mean ± SEM. Statistical analysis was performed using one-way ANOVA in A and C or two-way ANOVA in **E**, **G**, and **H** followed by Dunnett’s post hoc test. *p < 0.05* statistically significant. *\*\*\*P < 0.001* versus control; *^#^P < 0.05, ^##^P < 0.01, ^###^P < 0.001* versus control siRNA; *^@^P < 0.05*, *^@@^P < 0.01* versus Nrf2 siRNA; *^$$$^P < 0.001* versus injury in **A**. ‘ns’ is not significant.

*In vivo*, CD68^+^ macrophage infiltration into the perivascular space was increased following injury in WT mice and further enhanced in *Nrf2^−^/^−^* mice, whereas *ICAM-1^−^/^−^* mice exhibited minimal infiltration (Supplemental Fig. 2A and B).

To assess the roles of Nrf2 and ICAM-1 in post-TBI immune cell infiltration, we infused Calcein-AM-labeled monocytes or GFP-derived monocytes into WT, *Nrf2^−^/^−^* and *ICAM-17"* mice 24 h after injury. Fluorescence analysis revealed increased numbers of GFP^+^/Mac-1 ^+^ monocytes in *Nrf2^−^/^−^* mice, minimal GFP^+^/Mac-1^+^ cells in *ICAM-1^−^/^−^* mice, and intermediate levels in WT. These results indicate that Nrf2 limits, whereas ICAM-1 promotes, macrophage transmigration across the injured BBB (Fig. 7D-H).

### Nrf2 deficiency enhances ICAM-1–dependent neutrophil extracellular trap formation

Immunofluorescence analyses revealed robust induction of neutrophil extracellular traps (NETs), identified by co-localization of citrullinated histone H3 (H3Cit) and lymphocyte antigen 6 family member G (Ly6G), in the pericontusional cortex following TBI in WT mice. NET formation was significantly exacerbated in *Nrf2^−^/^−^* mice, both in terms of NET-positive area and the number of H3Cit7Ly6G^+^ structures, indicating that loss of Nrf2 markedly enhances neutrophil activation and chromatin extrusion after injury. In contrast, NET formation was nearly abolished in *ICAM-1^−^/^−^* mice, demonstrating that endothelial ICAM-1-mediated adhesion and transmigration are prerequisite events for NET release in the injured brain (Fig. 8A-C).

**Figure 8.**
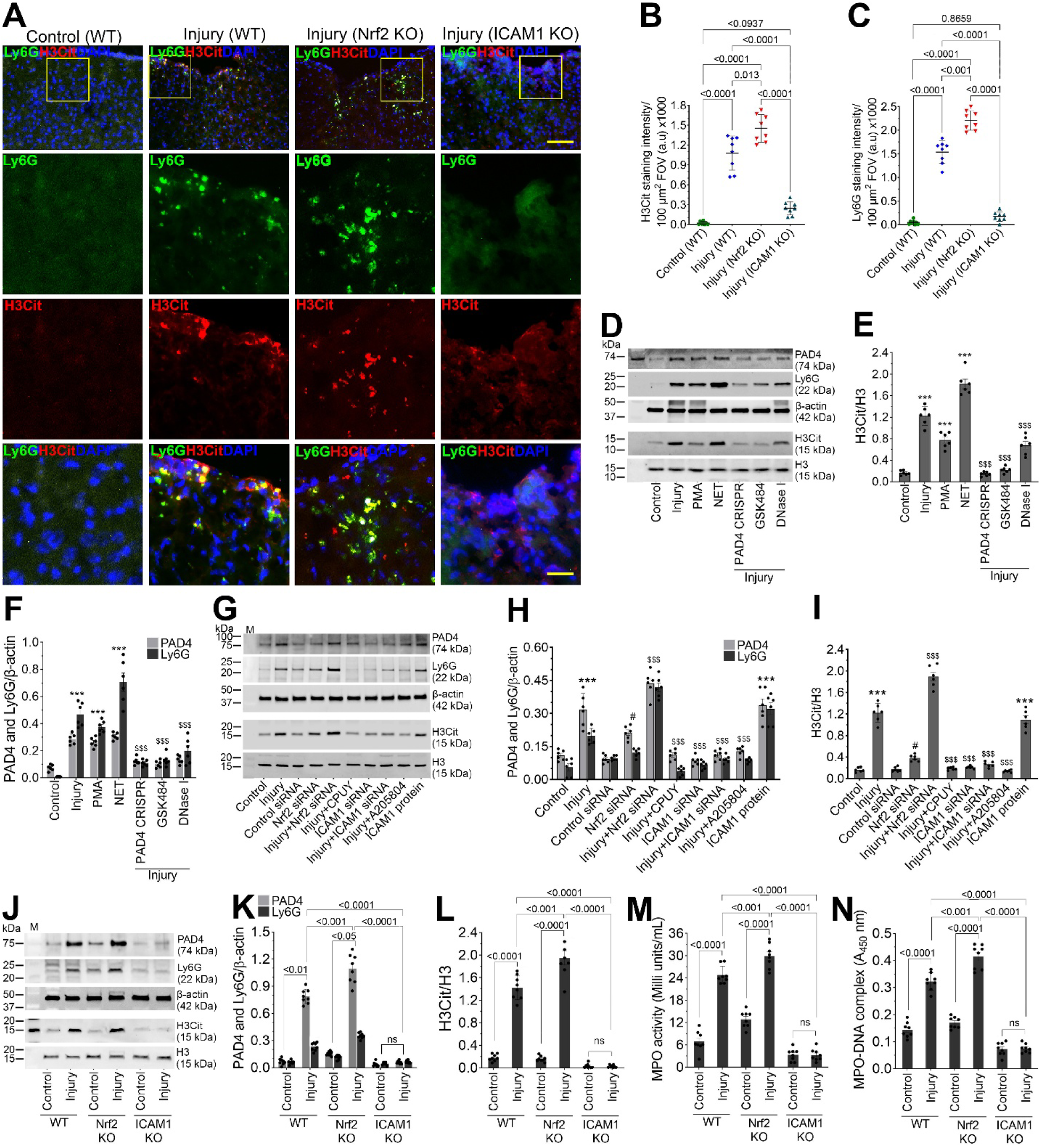
Nrf2 regulates neutrophil extracellular trap (NET) formation after traumatic brain injury. **(A-C)** Immunofluorescence analysis of NET formation in the pericontusional cortex of wild-type (WT), *Nrf2^−^/^−^*, and *ICAM-1^−^/^−^* mice following TBI. NETs were identified by co-localization of citrullinated histone H3 (H3Cit, a specific NET marker; red) and Ly6G (neutrophil marker; green). The nuclei were counterstained with DAPI (blue). Scale bar = 250 µm in the panels of first row, and 80 µm in the panels of second through fourth rows. Bar graphs show the quantification of H3Cit (B) and Ly6G (C) positive cells (n = 6/group). **(D-F)** Western blotting expression of PAD4, Ly6G, and H3Cit 24 hr after injury following genetic disruption of PAD4 (CRISPR/Cas9), pharmacological inhibition with GSK484, or DNase I, and the treatment with phorbol 12-myristate 13-acetate (PMA), or purified NETs in human brain microvascular endothelial cells (hBMVECs) exposed to injury. (**E** and **F**) Bar graphs represent densitometric quantification of H3Cit normalized to total H3 (E), and PAD4 and Ly6G normalized to β-actin (F) (n = 6/group). **(G-I)** Western blotting analysis of PAD4, Ly6G, and histone H3 citrullination in injured hBMVECs under Nrf2-deficient conditions and following inhibition of NET formation or ICAM-1 blockade. (**H** and **I**) Bar graphs represent densitometric quantification of PAD4 and Ly6G normalized to β-actin (H) and H3Cit normalized to total H3 (I) (n = 6/group). **(J-L)** Western blotting analysis of PAD4, Ly6G, and H3Cit expression in brain tissue from WT, *Nrf2^−^/^−^* and *ICAM-1^−^/^−^* mice under control and TBI conditions. (**K** and **L**) Bar graphs represent densitometric quantification of PAD4 and Ly6G normalized to β-actin (H) and H3Cit normalized to total H3 (I) (n = 8/group). **(M-N)** Measurement of NET-associated myeloperoxidase (MPO) activity (m) and MPO–DNA complexes (N) in brain tissue lysates from WT, *Nrf2^−^/^−^*, and *ICAM-1^−^/^−^* mice following TBI (n= 8/group). All values are expressed as mean ± SEM. Statistical analysis was performed using one-way ANOVA in **A**, **B**, **E**, **F**, **H**, and **I** and two-way ANOVA in **K-N**, followed by Dunnett’s post hoc test. *p < 0.05* statistically significant. *\*\*\*P < 0.001* versus control; *^#^P < 0.05, ^#^P < 0.05* versus control siRNA; *^$$$^P < 0.001* versus injury in **E**, **F**, **H**, and **I**. ‘ns’ is not significant.

We investigated NET formation in hBMVECs exposed to injury, PMA, or purified NETs. Injury and NET exposure increased PAD4, Ly6G, and H3Cit expression, with NETs eliciting the strongest response. PAD4 knockout using CRISPR/Cas9 PAD4, GSK484, or DNase I treatment attenuated these responses, confirming their specificity for NET-associated endothelial activation (Fig. 8D-F). Consistent with these findings, biochemical analyses demonstrated marked activation of NET-associated effector pathways. Injured hBMVECs exhibited significantly elevated MPO activity and increased formation of MPO-DNA and H3Cit-DNA complexes compared with controls. PMA or purified NET treatment reproduced these effects, whereas NET inhibition markedly reduced MPO activity and NET complex formation (Supplemental Fig. 3A-C). These data indicate that endothelial injury promotes robust NET-associated biochemical signaling that can be pharmacologically and genetically suppressed.

Upregulation of PAD4, and Ly6G, and increased histone H3 citrullination were significantly amplified by Nrf2 deficiency and effectively suppressed by pharmacological inhibition of NET formation or genetic disruption of PAD4, confirming a mechanistic link between oxidative stress, PAD4 activation, and NET generation. Notably, ICAM-1 inhibition markedly attenuated injury-induced PAD4 activation and NET marker expression, further supporting the requirement of ICAM-1-dependent neutrophil–endothelial interactions for NET release (Fig. 8G- I). Injury increased MPO activity and MPO–DNA and H3Cit–DNA complex levels, indicative of NET formation. These responses were amplified by Nrf2 deficiency and suppressed by PAD4 inhibition or NET blockade. ICAM-1 inhibition markedly reduced MPO activity and NET-associated DNA complexes, demonstrating its requirement for injury-induced NET release (Supplemental Fig. 4A-C).

Immunoblot analysis demonstrated injury-induced NET activation *in vivo*. WT mice showed increased PAD4 and Ly6G expression and elevated histone H3 citrullination following TBI, which were further amplified in Nrf2 knockout mice. In contrast, ICAM1-deficient mice exhibited minimal PAD4, Ly6G, and H3Cit expression under both control and injury conditions, indicating impaired neutrophil recruitment and NET formation (Fig. 8J-L).

Quantification of circulating and tissue NET markers corroborated these findings. ELISA-based measurement of H3Cit–DNA complexes revealed significant injury-induced increases in WT mice, which were further augmented in Nrf2 KO animals, both in brain tissue and plasma. In contrast, ICAM1 KO mice displayed persistently low H3Cit–DNA levels with no injury-associated elevation, demonstrating that ICAM1 is required for both local and systemic NET formation following TBI (Supplemental Fig. 5A-C).

Assessment of MPO activity in brain tissue lysate revealed a similar pattern. Injured WT mice showed a significant increase in MPO activity compared with controls, which was markedly amplified in Nrf2 KO mice. ICAM1 KO mice exhibited minimal MPO activity under both basal and injury conditions. ELISA-based quantification of MPO–DNA complexes confirmed robust injury-induced NET-associated MPO release in WT mice, further exacerbated by Nrf2 deficiency, whereas ICAM1 deletion effectively suppressed MPO–DNA complex formation (Fig. 8M and N). These trends were recapitulated in blood plasma too (Supplemental Fig. 6A and B).

Collectively, these data demonstrate that TBI induces robust neutrophil activation and NET formation that are potentiated by Nrf2 deficiency but critically depend on ICAM1-mediated neutrophil recruitment and adhesion. Nrf2 therefore acts as a key negative regulator of NET-driven neurovascular inflammation following traumatic brain injury.

### Nrf2 modulates ICAM-1–dependent vascular inflammation and attenuates neurological and cognitive deficits following TBI

Next, we analyzed motor coordination, spatial learning, memory, and anxiety-like behavior in WT, *Nrf2^−^/^−^*, and *ICAM-1^−^/^−^* mice following injury to determine the contribution of these genes to post-injury behavioral deficits. Motor coordination and balance were assessed using the rotarod test (Fig. 9A). Injured WT mice exhibited a significant reduction in latency to fall compared to uninjured controls, indicating impaired motor performance. This deficit was further exacerbated in *Nrf2^−^/^−^* mice, demonstrating greater motor dysfunction. In contrast, *ICAM-1^−^/^−^* mice showed significantly improved latency relative to injured WT animals, suggesting partial preservation of motor coordination.

**Figure 9.**
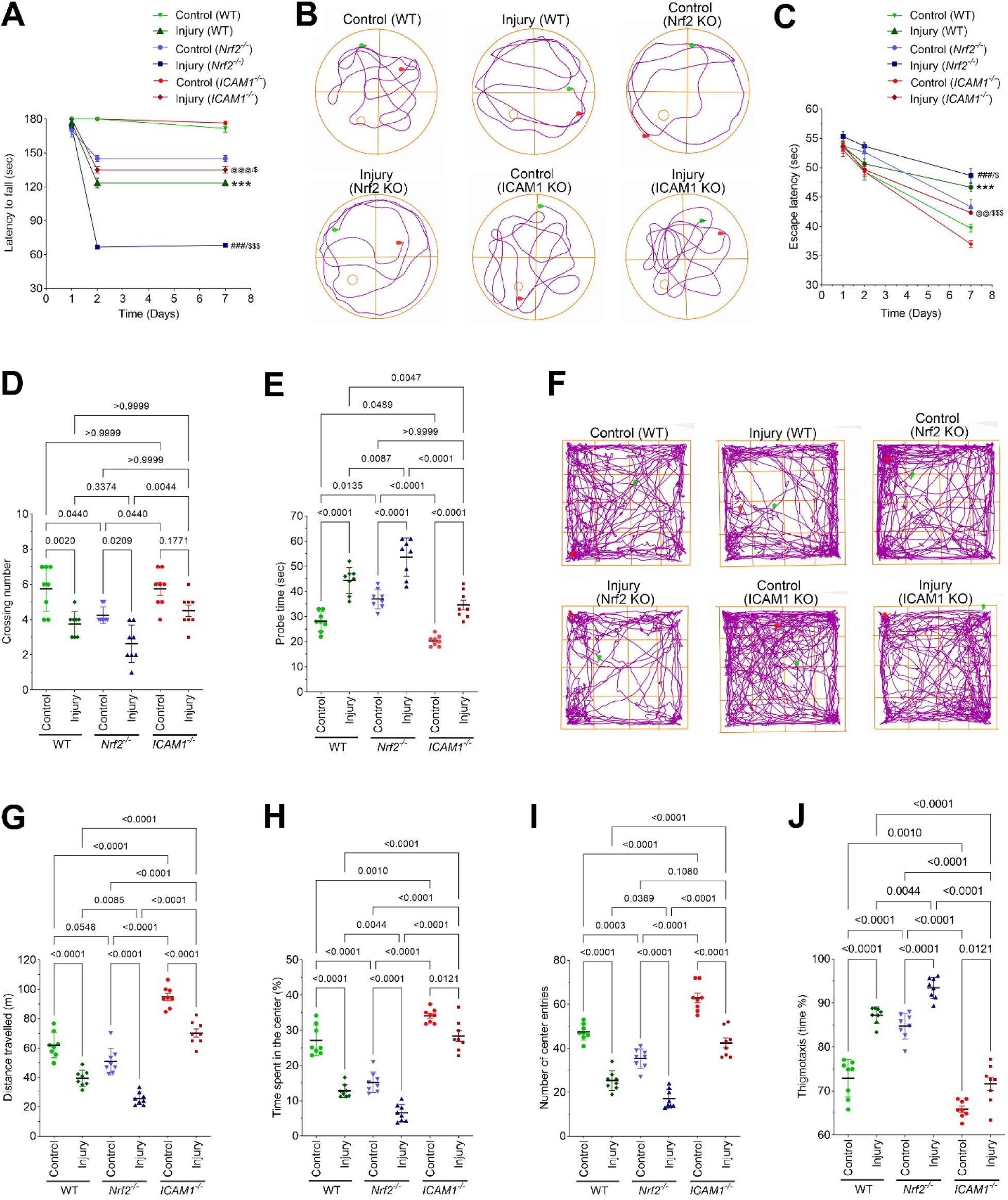
Nrf2 and ICAM-1 regulate motor, cognitive, and anxiety-like behavioral outcomes following injury. **(A)** Rotarod test assessing motor coordination and balance in wild-type (WT), *Nrf2^−^/^−^*, and *ICAM-1^−^/^−^* mice under control and injury conditions. Latency to fall was measured to evaluate motor performance (n = 8/group). **(B)** Representative swim trajectories from the Morris water maze showing search patterns during the probe trial in WT, *Nrf2^−^/^−^* and *ICAM-1^−^/^−^* mice following injury. **(C)** Escape latency during the acquisition phase of the Morris water maze test assessing spatial learning (n = 8/group). **(D-E)** Probe trial analysis of spatial memory, including the number of platform crossings (D) and time spent in the target quadrant (E) (n = 8/group). **(F)** Representative open-field movement tracks illustrating exploratory behavior patterns in WT, *Nrf2^−^/^−^* and *ICAM-1^−^/^−^* mice. **(G-J)** Quantification of open-field parameters including total distance traveled (G), time spent in the center (H), number of center entries (I), and thigmotaxis (J) (n = 8/group). All values are expressed as mean ± SEM. Statistical analysis was performed using two-way ANOVA followed by Dunnett’s post hoc test. *p < 0.05* statistically significant. *\*\*\*P < 0.001* versus control (WT); *^###^P < 0.001* versus control (*Nrf2^−^/^−^*); *^@@@^P < 0.001* versus control (*ICAM-1^−^/^−^*. *^$^P < 0.05, ^$$$^P < 0.001* versus injury (WT) in **A** and **C**.

Spatial learning and memory were evaluated using the Morris water maze (Fig. 9B-E). Representative swim trajectories revealed disorganized, peripheral search strategies in injured WT and *Nrf2^−^/^−^* mice, with relative preservation in *ICAM-17∼ mice* (Fig. 9B). During acquisition, injured WT mice showed increased escape latency compared to controls, a deficit that was significantly worsened in *Nrf2^−^/^−^* mice and attenuated in *CAM-1^−^/^−^* mice (Fig. 9C). In the probe trial, injured WT mice displayed reduced platform crossings and target quadrant time, impairments that were further aggravated in *Nrf2^−^/^−^* mice and significantly rescued in *ICAM-17"* mice (Fig. 9D and E).

Anxiety-like behavior and exploratory activity were assessed using the open-field test (Fig. 9F-J). Injured WT mice exhibited reduced total distance traveled, decreased center time and center entries, and increased thigmotaxis, indicating reduced locomotion and heightened anxiety-like behavior (Fig. 9F-J). These abnormalities were more pronounced in *Nrf2^−^/^−^* mice. Conversely, *ICAM-1^−^/^−^* mice demonstrated improved locomotor activity, increased center exploration, and reduced thigmotaxis compared to injured WT mice, consistent with attenuation of injury-induced anxiety-like behavior. Collectively, these data identify Nrf2 as protective and ICAM1 as a mediator of motor, cognitive, and anxiety-like dysfunction following injury.

## DISCUSSION

Traumatic brain injury (TBI) initiates a cascade of secondary injury processes that extend far beyond the initial mechanical insult, prominently involving oxidative stress, neuroinflammation, blood–brain barrier (BBB) dysfunction, and immune cell infiltration (1, 6, 26, 27, 40, 41). Although neuronal mechanisms of injury have been extensively studied, the molecular pathways governing neurovascular dysfunction remain incompletely understood. Here, we identify nuclear factor erythroid 2–related factor 2 (Nrf2) as a central regulator of endothelial homeostasis following TBI and demonstrate that impairment of Nrf2 signaling amplifies oxidative stress, ICAM-1–dependent leukocyte adhesion, matrix metalloproteinase activation, BBB disruption, immune cell transmigration, and neutrophil extracellular trap (NET) formation. These findings position Nrf2 not merely as a cytoprotective transcription factor, but as a master integrator of neurovascular defense following TBI.

A major finding of this work is that TBI leads to a marked impairment of Nrf2 signaling within the cerebral microvasculature. Under physiological conditions, Nrf2 activation is tightly regulated through Keap1-dependent cytoplasmic sequestration and stress-induced nuclear translocation, enabling transcription of antioxidant and cytoprotective genes (8–10, 42, 43). Despite evidence of endothelial nuclear localization after injury, both total and phosphorylated Nrf2 were significantly reduced in brain microvessels and in mechanically injured human brain microvascular endothelial cells, indicating that biomechanical trauma directly compromises endothelial Nrf2 signaling. These findings extend our and other labs’ previous reports of neuronal Nrf2 impairment after TBI (44, 45) and suggest that loss of vascular Nrf2 represents a critical but underappreciated contributor to secondary injury progression.

Consistent with this impairment, loss of Nrf2 activity resulted in a pronounced suppression of endothelial antioxidant defenses, including GPx1, GSTm1, HO-1, and NQO1. Restoration of these defenses through pharmacological disruption of the Keap1–Nrf2 interaction underscores the transcriptional dependence of vascular antioxidant capacity on intact Nrf2 signaling. ChIP-qPCR evidence further supports this conclusion by demonstrating reduced Nrf2 occupancy at antioxidant response elements following injury. Together, these findings suggest that diminished endothelial antioxidant reserve after TBI is not merely a consequence of oxidative stress but reflects a failure of adaptive redox transcriptional programs (44, 46, 47).

Failure to maintain antioxidant defenses translated into excessive oxidative and nitrosative stress within the neurovascular unit. Injury induced robust upregulation of NADPH oxidase 1 (NOX1) and inducible nitric oxide synthase (iNOS), along with accumulation of oxidative damage markers such as 4-hydroxynonenal and 3-nitrotyrosine (17, 48, 49). These results highlight the essential role of endothelial Nrf2 in maintaining vascular redox homeostasis and suggest that loss of this control creates a permissive environment for downstream inflammatory and proteolytic signaling (48, 50).One of the key mechanistic insights of this study is the identification of ICAM-1 as a critical downstream effector of Nrf2 dysfunction. Oxidative stress–driven ICAM-1 induction emerged as a central node linking endothelial Nrf2 suppression to leukocyte adhesion and transmigration. Pharmacological activation of Nrf2 markedly suppressed ICAM-1 expression, whereas genetic Nrf2 deficiency enhanced ICAM-1 levels even under basal conditions, indicating that Nrf2 actively restrains endothelial activation rather than passively responding to injury. These findings are consistent with redox-sensitive regulation of adhesion molecules and place Nrf2 upstream of vascular inflammatory signaling after TBI (20, 51).

ICAM-1 induction was functionally coupled to enhanced activation of leukocyte integrins LFA-1 and Mac-1, which are critical mediators of leukocyte adhesion and transmigration (25, 27), establishing a mechanistic basis for increased leukocyte–endothelial interactions. Genetic deletion of ICAM-1 abolished integrin activation, confirming its role as a pivotal mediator linking endothelial redox imbalance to immune cell recruitment. These findings are consistent with previous reports implicating ICAM-1 signaling in post-traumatic leukocyte infiltration and vascular inflammation (25, 27, 52).

The downstream consequences of ICAM-1 activation were evident at the level of endothelial junctional integrity. Tight junction proteins, including occludin, claudin-5, and ZO-1, were profoundly disrupted following injury, with further exacerbation under Nrf2-deficient conditions. These findings suggest that Nrf2 protects BBB integrity not only by limiting oxidative stress but also by suppressing ICAM-1–dependent pathways that compromise endothelial junctions (14, 48, 53). Beyond tight junctions, adherens and gap junction proteins critical for endothelial–pericyte communication, such as N-cadherin and connexin-43, were similarly compromised. Restoration of these junctional proteins by Nrf2 activation or ICAM-1 inhibition underscores the importance of Nrf2-dependent suppression of inflammatory signaling in preserving BBB architecture.

Functional assays confirmed that these molecular changes translated into significant BBB dysfunction. Increased permeability to tracers in vitro and in vivo, along with elevated circulating markers of BBB breakdown and neuronal injury, establish a causal relationship between Nrf2 impairment, ICAM-1 activation, junctional destabilization, and barrier failure. These findings reinforce the concept that endothelial redox regulation is a determinant of BBB integrity after TBI.

Beyond leukocyte recruitment, this study reveals a previously unrecognized role for endothelial Nrf2 in regulating neutrophil extracellular trap formation. NET generation was markedly increased following TBI and further amplified by Nrf2 deficiency, whereas ICAM-1 deletion nearly abolished NET formation. These data suggest that endothelial oxidative stress and adhesion signaling are prerequisites for NET release (27) and position Nrf2 as a negative regulator of immunothrombotic responses within the injured brain. Given the emerging role of NETs in vascular injury, thrombosis, and neuroinflammation (29, 31, 34, 54, 55), this finding substantially broadens the pathophysiological relevance of endothelial Nrf2 signaling.

A major strength of this study is its multilevel mechanistic analysis of Nrf2 signaling within the neurovascular unit after TBI, integrating in vivo genetic models, in vitro endothelial injury paradigms, and molecular, biochemical, and functional BBB assays. The combined use of WT, *Nrf2^−^/^−^*, and *ICAM-1^−^/^−^* mice together with pharmacological and genetic manipulation of hBMVECs establishes clear causal links between endothelial Nrf2 suppression, oxidative stress, inflammatory activation, BBB disruption, and immune cell infiltration. Another key strength is the strong endothelial specificity of the findings. Microvessel-enriched analyses, vessel-restricted immunofluorescence, endothelial stretch-injury models, and ChIP-qPCR evidence of reduced Nrf2 binding to antioxidant gene promoters collectively demonstrate that Nrf2 dysfunction occurs directly within the cerebral microvasculature. The study further excels in linking redox imbalance to inflammatory and proteolytic cascades, identifying an Nrf2–ICAM-1 axis that drives leukocyte adhesion, junctional destabilization, and BBB permeability. Notably, the identification of Nrf2 as a negative regulator of NET formation after TBI provides a novel mechanistic connection between endothelial oxidative stress, neutrophil activation, and NET formation. Finally, the consistency of findings across cell culture systems, animal models, imaging, biochemical assays, permeability measurements, and circulating biomarkers adds substantial rigor and internal validity to the conclusions.

Our behavioral analyses demonstrate that Nrf2 and ICAM1 play opposing roles in post-injury neurological function. Nrf2 deficiency worsened motor coordination, spatial learning, memory, and anxiety-like behavior, highlighting its protective role in maintaining neuronal and cognitive integrity after injury. In contrast, ICAM1 deficiency mitigated these deficits, suggesting that ICAM1 contributes to injury-induced functional impairment. These results are consistent with previous evidence linking Nrf2 to antioxidant defense and neuroprotection, and ICAM1 to inflammation-mediated neuronal dysfunction, supporting a model in which modulation of these pathways can influence recovery of motor and cognitive performance following brain injury.

Several limitations should be acknowledged. Some biochemical analyses relied on whole cortical lysates, which may include non-endothelial contributions. Future studies using endothelial-specific Nrf2 conditional knockout models will further refine cell-type resolution. In addition, the study focused on acute and subacute time points; long-term effects of sustained Nrf2 suppression on chronic vascular remodeling and neurodegeneration remain to be determined. While pharmacological Nrf2 activation was effective, off-target effects of systemic activation cannot be fully excluded, highlighting the need for endothelial-targeted or peptide-based delivery approaches.

## Conclusions

In summary, this study identifies Nrf2 as a master regulator of neurovascular protection following traumatic brain injury. Impairment of Nrf2 signaling initiates a cascade of pathological events involving oxidative stress, ICAM-1 activation, leukocyte adhesion, junctional protein disruption, BBB breakdown, immune cell infiltration, and NET formation. Consistently, Nrf2 deficiency exacerbated, whereas ICAM-1 deficiency attenuated, motor deficits, spatial learning and memory impairments, and anxiety-like behavior, linking neurovascular dysfunction to functional outcomes. Therapeutic strategies aimed at restoring Nrf2 activity or targeting ICAM- 1–dependent pathways may therefore offer a multifaceted approach to mitigating secondary neurovascular injury and improving outcomes after TBI.

## MATERIALS AND METHODS

### Sex as a biological variable

Both male and female mice were included in all in vivo experiments to account for sex as a biological variable. Animals of both sexes were randomly assigned to experimental groups, and sex was balanced across treatment and genotype groups to the extent possible. The present study was designed to determine the overall effects of traumatic brain injury, Nrf2 deficiency, and ICAM-1 deficiency on neurovascular inflammation, leukocyte recruitment, and NET formation and was not powered to detect sex-specific differences or interactions between sex and experimental conditions. Therefore, data from male and female mice were analyzed together. We did not observe an obvious sex-dependent pattern in the measured outcomes; however, formal sex-stratified statistical comparisons were not performed. Inclusion of both sexes increases the generalizability of the findings across biological sex.

### Animals and fluid Percussion Injury

Male and female C57/BL6 mice wild-type (WT), Nrf2 knockout (*Nrf2^−^/^−^)* mice and ICAM-1 knockout (*ICAM-1^−^/^−^)* mice (9 weeks old, 20-25 g; Jackson Laboratory, Bar Harbor, ME) were used for this study. Animals were housed under temperature-controlled conditions with a 12-h light/dark cycle and provided ad libitum access to food and water. Lateral fluid percussion injury (FPI) and sham procedures were performed according to our previously established protocols (1, 7, 27, 45, 56). Briefly, mice were anesthetized with a ketamine/xylazine mixture (80 mg/kg ketamine and 10 mg/kg xylazine, i.p.) and surgically implanted with a Luer-Lok syringe hub on the right side of the skull using a stereotaxic device, approximately 24 h prior to injury. The hub surrounded a craniotomy of identical size positioned 3.0 mm posterior and 3.5 mm lateral to the bregma. An additional cap was placed around the syringe hub, and two screws were implanted into the skull to provide additional stability. Cranioplastic dental cement (AM Systems, Carlsborg, WA) was applied around the syringe hub and between the cap to ensure proper fluid transmission and structural support.

Immediately prior to injury, mice were anesthetized with 5% isoflurane until the foot-pinch reflex was absent. Animals were then connected to a digitally controlled FPI system (FP302; AmScien Instruments, Richmond, VA), and injury was induced at 25 psi (moderate injury) with a pressure rise time of 8 ms. Following injury, mice exhibited apnea, loss of consciousness (LOC), and hyperextension of the tail and hind limbs. Sham control mice underwent identical anesthesia and surgical procedures and were connected to the FPI device but did not receive injury.

A total of eight mice per experimental group were used. Based on our previous studies and power analyses, six animals per group are sufficient to achieve statistically significant outcomes (1, 7, 27, 40). At 24 h post-injury, animals were anesthetized with a ketamine/xylazine mixture and transcardially perfused with 1x PBS followed by 4% paraformaldehyde. Brains were harvested, embedded in optimal cutting temperature (OCT) compound, and stored frozen until further analysis.

### Cell culture

hBMVEC (ScienCell Research Laboratories (Carlsbad, CA) will be cultured in 6-well Bioflex culture silicone membrane plates. Once a monolayer of hBMVECs is formed, a small number (5,000 cells/per well) of human neutrophils (Stemcell Technologies, Cambridge, MA) will be added and co-cultured for 48 hr. The addition of neutrophils will induce the formation of NET in hBMVEC culture(57). For experiments with pharmacological treatments and gene manipulations, the cells will be treated with CP (5 µg/mL), PAP (5 µg/mL), GSK484 (a PAD4 inhibitor, 10 µM)(58, 59), rPAD4 (PAD4 recombinant protein, 2.5 µg/mL), PMA (50 nM; Sigma Aldrich), or DNase-I (100 U/mL; Worthington Biochem) 30 min prior to stretch injury. Then the cells will be exposed to stretch injury (biaxial) with a pressure of 3.0 psi (moderate injury for the cells(4)) as we reported(3, 5, 60) using a biaxial *in vitro* cell injury device- “cell injury controller II” (Custom design and fabrication Inc., Sandstom, VA) as we and others described previously(4, 40, 45, 60–62). Control cells will not be subjected to stretch injury. PMA is an activator of protein kinase C (PKC) that can trigger NET formation(63). DNase-I has a role in disrupting the formation of NET(64). Since GSK-484 cannot be used for clinical development(65) (see innovation section), we propose to use it *in vitro* studies only to show the role of PAD4 in NET formation. The experimental groups are given in Figure 3A.

Human brain microvascular endothelial cells (hBMVECs; Cell Biologics, Chicago, IL) were used between passages 3-4, with no observable passage-dependent differences. Cells were cultured in DMEM/F-12 medium (Gibco, CA, USA) supplemented with 10 mM HEPES, 13 mM sodium bicarbonate (pH 7.0), 10 mmol/L L-glutamine, 10% fetal bovine serum (FBS; Atlanta Biologicals), penicillin and streptomycin (100 µg/mL each; Thermo Fisher Scientific), 1% heparin (Thermo Fisher Scientific), 1% endothelial cell growth supplement (ECGS; BD Biosciences), and 1% amphotericin B (Fungizone; Caisson Labs). Cells were maintained at 37 °C in a humidified atmosphere of 5% CO_2_ and seeded onto BioFlex 6-well culture plates (Flexcell International Corp.) pre-coated with type I rat-tail collagen (0.1 mg/mL; Gibco) diluted in sterile double-distilled water. Once a monolayer of hBMVECs is formed, a small number (5,000 cells/per well) of human neutrophils (Stemcell Technologies, Cambridge, MA) will be added and co-cultured for 48 hr. The addition of neutrophils will induce the formation of NET in hBMVEC culture (57). Culture medium was replaced every 2 days until tight endothelial monolayers formed, typically within 6–8 days.

### *In vitro* stretch injury and treatments

Following the formation of tight monolayers (approximately day 7), hBMVECs and human neutrophils co-culture grown on BioFlex plates were subjected to biaxial stretch injury at ∼3.0 psi, as described previously (1, 27). After stretch injury, cultures were returned to the incubator and maintained for the indicated experimental duration (24 h). For pharmacological and genetic manipulation studies, cells were treated 30 min prior to stretch injury with Nrf2 siRNA (Cat. No. SC37049; Santa Cruz Biotechnology), ICAM-1 siRNA (Cat. No. SC29354; Santa Cruz Biotechnology), scrambled control siRNA (Cat. No. SC37007; Santa Cruz Biotechnology), CPUY192018 (10 µM; an inhibitor of the Keap1–Nrf2 protein interaction; Cat. No. AOB9974; AOBIOUS), or A205804 (100 ng/mL; ICAM-1 inhibitor; Cat. No. 21252; Cayman Chemical). At the designated time points post-injury, cells were lysed for protein extraction, culture supernatants were collected for cytokine analysis, and cells were fixed for immunostaining.

### PAD4 CRISPR/Cas9 treatments

We validated the role of peptidyl arginine deiminase 4 (PAD4) in NET formation (66, 67) by knocking down the PAD4 gene using CRISPR/Cas9 technology. A PAD4 CRISPR All-in-One AAV vector containing saCas9 (mouse; serotype 9; pAAV-PGK-saCas9-U6-sgRNAsa) and a scrambled CRISPR/Cas9 AAV control vector (Control CRISPR; Cat. No. K079), both driven by a PGK-U6 promoter, were purchased from Applied Biological Materials Inc. (Richmond, BC, Canada). Adeno-associated virus (AAV), a parvoviral vector classified under Risk Group 1 (not associated with disease in healthy adult humans (68) was shipped frozen on dry ice. Upon receipt, viral stocks were stored at -80 °C for long-term storage, aliquoted into single-use siliconized screw-cap microcentrifuge tubes, and stored at -20 °C. All AAV handling was performed under biosafety level-1 (BSL-1) containment in accordance with NIH and OSHA guidelines. Surfaces and materials exposed to AAV were disinfected using 0.5% sodium hypochlorite, 2% glutaraldehyde, or autoclaving for 30 min.

For gene knockdown experiments, hBMVEC–neutrophil (Cat. No: 200-0384, Stemcell Technologies) co-cultures were treated with CRISPR AAV at a concentration of 1.1 x 10d genome copies (GC)/mL according to the manufacturer’s instructions. CRISPR treatment was administered immediately following injury. At 24 h post-injury, cell lysates and culture supernatants were collected for mRNA, protein, and cytokine analyses. The cytotoxicity of CRISPR/Cas9 AAV treatment was assessed using an *MTT assay* (*69*). No significant cytotoxicity or adverse effects on cell viability were observed following PAD4 deletion under basal conditions.

### Analysis of ROS and NO

Quantification of reactive oxygen species (ROS) and nitric oxide (NO) (n = 6/group) was performed using the CellROX Green Reagent Kit (Cat. No. C10444) and the DAF-FM diacetate Kit (Cat. No. D-23844) (Thermo Fisher Scientific, Rockford, IL), respectively, in hBMVECs 24 h following stretch injury, according to the manufacturer’s instructions (4, 70). For oxidative stress analysis, 24 h after stretch injury, with or without drug/inhibitor treatment, CellROX Green Reagent was added to the cells at a final concentration of 5 µM and incubated for 30 min at 37 °C. Cells were then washed three times with 1x PBS, fixed with a formaldehyde-based fixative (4% paraformaldehyde for 15 min), and fluorescence signals were analyzed within 24 h. For NO quantification, 24 h after stretch injury and drug/inhibitor treatment, cells were incubated with DAF-FM diacetate (1-10 µM) for 20-60 min at 4 °C-37 °C. The medium was then replaced with fresh culture medium and cells were incubated for an additional 15-30 min to allow complete de-esterification of the intracellular diacetates. Fluorescence was measured using excitation and emission wavelengths of 495 nm and 515 nm, respectively.

### Immunostaining and microscopy

**a) Immunofluorescence:** The cultured hBMVEC (on silicone membrane) and 10 pm thickness coronal brain tissue sections were washed with PBS and fixed in 4% paraformaldehyde for 20 minutes at 25°C. The washed membrane or tissue sections were then blocked with 3% normal goat serum containing 0.1% Triton X-100 at 25°C for 1 hr followed by overnight incubation with primary antibodies at the concentration of 3.0 µg/mL at 4°C. The cell culture membranes or tissue sections were incubated with primary antibody to anti Nrf2, anti-p- Nrf2, anti-ICAM-1, anti-LFA-1, anti-occludin, anti-claudin5, anti-ZO-1, anti-N-cadherin. To show the cell specificity of the expression, in double immunofluorescence, we incubated cells or tissue sections with sheep anti-vWF (endothelial specific marker). After washing with 1x PBS, the cells or tissue sections were incubated with Alexa Fluor 488 or 594 conjugated with anti-mouse or anti-rabbit immunoglobulin G (IgG) (1:500 dilution or 4 µg/mL) for 1 hr and mounted with 5-10 µL immuno mount containing DAPI (Invitrogen) on a slide. The silicone membrane was carefully cut at the edges using a sterile scalpel blade and mounted on a slide using DAPI (40). The photographs were captured using the Eclipse TE200 fluorescent microscope (Nikon, Melville, NY) and NIS Elements software (Nikon, Melville, NY) in three channels, DAPI (cell nuclei), Alexafluor-488 (green color emission), and Alexa fluor-594 (red color emission). After staining, the slides mounted with tissue sections were randomly chosen for quantification, which was done blindly by masking the slide label. For quantitative analysis, we used at least 4 tissue samples for immunofluorescent staining and captured 6 images from a single sample (slide). Just below the injury or the peri-lesion area, approximately 1.5 mm^2^ areas were covered for the analysis. For consistency, we kept the same parameters of camera and software including the brightness of the excitation light, the detector sensitivity (gain), or the camera exposure time across the samples or tissue sections. The intensity of immunostaining was analyzed by ImageJ (NIH) software. To determine the percentage of positive cells for a particular protein, we threshold the images by keeping lower threshold level 80 and upper threshold level 200. We kept a uniformly dark background for all images for correcting uneven illumination in fluorescence images.
**b) HRP-substrate immunohistochemistry**: Coronal brain sections (10 µm thick) fixed with paraformaldehyde were washed in 1x PBS and incubated in 0.3% H_2_O_2_ and 0.3% Triton X-100 in 1x PBS for 20 min at room temperature. Tissue sections were then rinsed three times with 1x PBS and blocked with 2% normal goat serum in phosphate buffer containing 0.3% Triton X-100 for 20 min at room temperature. After three washes with 1x PBS, sections were incubated overnight at 4 °C with rabbit anti-ICAM-1 primary antibody. The primary antibody was removed by washing three times with 1× PBS, followed by incubation with HRP-conjugated goat secondary antibody (1:200 dilution) in PBS containing 0.3% Triton X-100 for 1 h at room temperature. Sections were then washed three times with 1x PBS. Peroxidase substrate reagent (SK-4600; Vector Laboratories) containing 0.02% H_2_O_2_ was applied to the tissue sections and incubated for 5-10 min until color development was observed under a dissecting microscope. The reaction was stopped by washing the sections three times with 1x PBS for 5 min each. Finally, tissue sections were mounted using Histomount and glass coverslips for microscopic analysis.

### Western blotting

Protein lysates from *in vitro* and *in vivo* experiments were prepared as described previously (1, 27, 71). Briefly, cells (three wells from the same treatment group) and tissues (50 mg of cortical tissue collected below the injury site from the neocortex) were lysed using CellLytic-M buffer (Thermo Scientific) supplemented with a protease inhibitor cocktail (Sigma). Protein concentrations from both *in vitro* and *in vivo* homogenates, as well as cell culture supernatants, were determined using the bicinchoninic acid (BCA) assay (Thermo Scientific, Rockford, IL). Protein extracts (15 µg) were separated on 4-15% gradient SDS–PAGE gels (Bio-Rad, Hercules, CA), transferred onto nitrocellulose membranes, and probed with the appropriate primary antibodies (1 µg/mL; see Supplementary Table 1), followed by horseradish peroxidase–conjugated secondary antibodies (1:5000; Fisher Scientific). Immunoreactive bands were visualized using an enhanced chemiluminescence reagent (Advansta) and imaged with a Bio-Rad ChemiDoc Imaging System. Mouse anti-β-actin (1 pg/mL) was used as a loading control for normalization. Western blot bands were quantified as arbitrary densitometric intensity units using ImageJ software.

### Coimmunoprecipitation

Co-immunoprecipitation was performed to analyze ICAM-1, LFA-1, and Mac-1 protein–protein interactions using the Pierce™ Classic Magnetic IP/Co-IP Kit (Cat. No. 88804). Briefly, brain tissues (50 mg) from WT and *Nrf2^−^/^−^* mice, with and without FPI, were harvested and lysed in 500 µL of IP lysis/wash Buffer. Following lysis and centrifugation, the supernatants were transferred to new tubes for protein concentration determination. For immunoprecipitation, Pierce protein A/G magnetic beads were used according to the manufacturer’s instructions. A total of 500 µg of protein from each sample was incubated with 10 µg of anti-ICAM-1 antibody overnight at 4°C. After immune complex formation, Protein A/G magnetic beads were added and incubated for 1 h at room temperature. Following washing of the immunoprecipitated complexes, the samples were resuspended in 200 µL of 2x SDS sample buffer. Aliquots (20 µL) were then subjected to Western blot analysis using antibodies against ICAM-1, LFA-1, and Mac-1.

### Reverse-transcription quantitative polymerase chain reaction (RT-qPCR)

The mRNA expression levels of Nrf2, glutathione peroxidase-1 (GPx1), glutathione S-transferase m1 (GSTm1), heme oxygenase-1 (HO-1), and NAD(P)H quinone oxidoreductase-1 (NQO1) were determined by reverse transcription-quantitative polymerase chain reaction (RT-qPCR). At 24 h post-TBI, WT and *Nrf2^−^/^−^* mice were euthanized, brains were removed, and tissues were stored at -80 °C. Total RNA from the injured cortex was extracted using the Qiagen RNeasy Mini Kit (Cat. No. 74104) according to the manufacturer’s instructions, followed by DNase I treatment for 20 min at 37 °C to eliminate genomic DNA contamination.

RNA quantity, purity, and integrity were assessed using Qubit 2.0 (Thermo Fisher Scientific, USA) and an Agilent Bioanalyzer (Agilent Technologies, USA). Isolated total RNA was reverse transcribed into cDNA using the iScript cDNA Synthesis Kit (Bio-Rad, USA). Primer sequences are listed in Supplementary Table 2, and PCR conditions are described below. Briefly, PCR amplification was performed using 25 ng of cDNA template mixed with gene-specific forward and reverse primers and iTaq Universal SYBR Green Supermix (Bio-Rad Laboratories, USA) in a 20 µL reaction volume on a StepOne Real-Time PCR System (Applied Biosystems, USA) under standard cycling conditions. GAPDH was used as the endogenous control, and relative mRNA expression levels were calculated using the AACt method.

### Chromatin immunoprecipitation-qPCR Assay (ChIP-qPCR)

Brain tissue was extracted from WT and *Nrf2^−^/^−^* mice and crosslinked with 1% formaldehyde at room temperature, and subsequently quenched with glycine. After washing with 1XPBS, the fixed samples were lysed in 500 pl cell lysis buffer (Cell lytic buffer, Sigma) and collected nuclei were stored at -80°C as nuclear pellets or nuclear lysates dissolved in nuclei lysis buffer (50-mM Tris-HCl pH8.0, 10-mM EDTA, 1% (wt/vol) SDS and protease inhibitor). The resulting nuclear lysate was sonicated until crosslinked chromatin was sheared to an average length of 0.3∼1.0 kb. Supernatant (5 pl) was used as an input control. The remaining lysate was diluted 10-fold with ChIP dilution buffer (16.7 mM Tris–HCl, 167 mM NaCl, 0.01% SDS, 1.1% Triton X-100, 1.2 mM EDTA, and protease inhibitor) and was incubated with Nrf2 antibody, followed by DNA/protein A-agarose magnetic beads. Bound protein-DNA complexes were eluted with a solution containing 0.1 M NaHCO_3_ and 1% SDS (elution buffer). After the cross-linking was reversed, chromatin fragments were treated with RNase A and proteinase K and DNA were purified with phenol-chloroform-isoamyl alcohol extraction. The DNA sample was then subjected to quantitative PCR using a StepOne Real-Time System (Applied Biosystems, USA) with standard cycling conditions. In the ChIP-qPCR analyses, the values from the immunoprecipitated samples were normalized to that from the input DNA. Primer sequences are provided in Supplementary Table 2.

### Luciferase reporter assay

Luciferase reporter assays were performed to assess Nrf2- dependent antioxidant response element (ARE) transcriptional activity of GSTm1, GPx1, HO-1, and NQO1 in endothelial cells. Endothelial cells were seeded in 24-well plates and transiently transfected with pGL3-GSTm1-Luc, pGL3-GPx1-Luc, pGL3-HO-1-Luc, or pGL3-NQO1-Luc promoter reporter plasmids containing functional ARE sequences, along with pRL-TK Renilla luciferase plasmid (Promega) as an internal control for transfection efficiency. Transfections were performed using a lipid-based transfection reagent according to the manufacturer’s instructions. Following transfection, cells were treated as indicated and incubated for 24-48 h. Firefly and Renilla luciferase activities were measured using a dual-luciferase reporter assay system using a Promega GloMax luminometer. Firefly luciferase activity was normalized to Renilla luciferase activity, and results were expressed as relative luciferase units (RLU) compared with control, reflecting Nrf2-mediated activation of ARE-driven gene transcription.

### *In vitro* monocyte adhesion assay

The adhesion of immune cells to the hBMVEC monolayer was performed *in vitro* as we and others described previously (72–76) with minor modifications. Briefly, hBMVEC were cultured on rat tail collagen (0.1 mg/mL) coated Bioflex 6-well culture plates at a density of 250,000 cells/well at 37°C in a dehumidified atmosphere of 5% CO2 (see cell culture section) until tight monolayer formed. The cells were treated with Nrf2 siRNA, ICAM- 1 siRNA, control siRNA, CPUY192018, or A205804 and after 30 min the stretch injury (3 psi) was conducted to the cells. 24 hours after the injury, Calcein-AM (a green fluorescence cell tracker; 2µM, Invitrogen)-labeled primary human monocytes were introduced into the endothelial monolayers (1X10^6^ monocytes/well) and the cells were then allowed to adhere for 2 h at 37°C in a tissue culture incubator. Non-adherent cells were removed by washing with 1x PBS. The relative fluorescence intensity of the adhered cells was detected by a fluorescence-based assay using the Tecan Genios fluorescence plate reader and spectrofluorometric quantification was performed at 492 nm (excitation) and 535 nm (emission) on a plate reader. The actual numbers of adhered monocytes were calculated from the internal standard curve of the labeled cells, and the data were presented as the fold difference of the untreated control.

### *In vitro* BBB permeability assay

To analyze the role of Nrf2 and ICAM-1 in BBB permeability in cells, the hMBECs (100,000 cells/insert) were cultured on type I collagen coated FluoroBlok tinted tissue culture inserts (with 3 µm pores, BD Biosciences) and after the formation of the tight monolayer, cell cultures were treated with siRNA and inhibitors of Nrf2 and ICAM-1 for 24 hr, then FITC-labelled Dextran-4 (5 µM, Invitrogen) were introduced into the endothelial monolayers. FITC-labelled Dextran-4 were then allowed to cross through the monolayers for 2 hr at 37°C in tissue culture incubator in the absence of test compounds. The relative fluorescence intensity of the crossed FITC-labelled Dextran-4 at the lower chamber and those FITC-labelled Dextran-4 that was stuck between the porous membranes and the lower chambers were detected by a fluorescence-based assay Tecan Genios fluorescence plate reader (excitation 492 nm and emission 535 nm). The actual quantity of monolayer crossed FITC-labelled Dextran-4 was calculated, and the data were presented as the fold difference of the untreated control cells.

### *In vivo* cell adhesion and migration assay

Femoral bones from euthanized mice were dissected out under sterile conditions and washed in 1X PBS. Bone marrow was flushed out repeatedly with 1X Hanks’ Balanced Salt Solution (HBSS) through the cut ends of the bones using a 1ml syringe. The bone marrow suspension was filtered through a 40 pm cell strainer. The filtrate was collected and centrifuged at 1, 800 rpm for 5 min at 4°C. The pellet was suspended in 1 ml of DMEM/F-12 media containing 10% fetal bovine serum (FBS), penicillin and streptomycin (100 pg/ml each, Invitrogen) and 0.001% MCSF (500 pL in 500 mL media). Then the cells were dissociated by trituration (10–15 times) and counted with Trypan Blue using Hemocytometer. Bone marrow-derived cells were differentiated to macrophages in culture containing the macrophage-colony stimulating factor (MCSF) in the culture medium (plating 2X10^6^ cells/T-75 flask). Cells were fed every third day. After 6 days, the differentiated macrophages were detached by a cell scraper, suspended in 1 ml HBSS and centrifuged at 1800 rpm for 5 min at 4°C. Cell pellets were then dissociated by repeated trituration, counted and were labeled with Calcein-AM (Molecular Probes, Invitrogen) in 1 ml of HBSS at 37°C for 15 min. Excess unbound Calcein-AM was washed out by centrifugation and cell pellets resuspended in HBSS (2X10^6^ cells) were infused through the common carotid artery using 30 G needle. The animal was euthanized 1 h after the cell infusion and brain microvessels were surgically removed for observation of calceinAM labeled macrophages under the fluorescent microscope.

To further validate transmigration, we infused GFP monocytes differentiated macrophages to the carotid artery of mice and analyzed the presence of GFP macrophages in the brain by immunostaining with anti-GFP antibody. Mac-1 marker was co-stained for confirming that the co-localization was in macrophages. Briefly, the GFP monocytes were separated from GFP transgenic mice (C57BL/6-Tg(CAG-EGFP)131Osb/LeySopJ; Jackson Laboratory) and cultured as per the protocol mentioned above. Bone marrow-derived cells were differentiated to macrophages in culture containing the macrophage-colony stimulating factor (MCSF) in the culture medium (plating 2X10^6^ cells/T-75 flask). After 6 days, the differentiated macrophages were detached by a cell scraper, suspended in 1 ml HBSS and centrifuged at 1800 rpm for 5 min at 4°C. Cell pellets were then dissociated and infused through the common carotid artery using 30 G needle. The animal was euthanized 1 h after the cell infusion and brain tissue sections were prepared for observation of GFP macrophages by immunofluorescence staining with anti-GFP antibody and co-stained with anti-Mac-1 antibody and visualized and captured pictures using the fluorescent microscope.

### *In vivo* BBB permeability assay

Assessment of BBB permeability in the brain following FPI was determined by intravascular infusion of sodium fluorescein (Na-Fl) and Evan’s Blue (EB) tracer and subsequent detection of extravasated molecules in the brain tissue as we previously described (6, 7, 76). In brief, after 24 h of FPI injury, Na-Fl/EB dye mixtures (5 pM each) were infused into the common carotid artery and two hours after the infusion, the animals were decapitated, and the brains were removed, dissected, weighed, and homogenized in 600 pl 7.5%(w/v) trichloroacetic acid (TCA). Resulting suspensions were divided into two 300 pl aliquots. One aliquot was neutralized with 50 pl of 5N NaOH and fluorescence was measured on a multifunctional Promega microplate reader (excitation 485nm, emission 535nm) to determine Na-Fl concentration. The second aliquot was centrifuged for 10 min at 10,000 rpm and 4^°^C, and the EB concentration in the supernatant was measured by absorbance spectroscopy at 620 nm. A standard curve was generated using serial dilutions of EB/Na-Fl solution in 7.5%TCA.

### Enzyme-linked immunosorbent assay (ELISA)

To quantify the NET markers, we analyzed the H3Cit-DNA complex and myeloperoxidase-DNA (MPO-DNA) complex formation brain tissue lysates, and blood plasma protein samples from control and injury WT, *Nrf2^−^/^−^* and *ICAM-1^−^/^−^* mice subjected to 25 psi FPI. Using commercial ELISA kits, the level of H3Cit-DNA and MPO-DNA were analyzed in tissue lysates as per manufacturer’s instructions.

### Behavioral analysis

Behavioral assessments were conducted to evaluate motor, cognitive, and anxiety-like functions in wild-type (WT), *Nrf2^−^/^−^*, and *ICAM-1^−^/^−^* mice under control and injury conditions (n = 8/group) (1, 27, 77–80). All behavioral tests were performed within the 12/12 h light/dark cycle, and experimenters were blinded to the treatment groups as described previously(1, 56). Motor coordination and balance were measured using the rotarod test, with latency to fall recorded. Spatial learning and memory were assessed using the Morris water maze, including acquisition trials to determine escape latency and probe trials to evaluate platform crossings and time spent in the target quadrant. Anxiety-like and exploratory behaviors were analyzed using the open-field test, with parameters including total distance traveled, time spent in the center, number of center entries, and thigmotaxis. Behavioral tracking and analysis were performed using standard automated video-tracking systems.

### Quantification and statistical analysis

Sample sizes were prospectively derived by conducting power analyses using G*Power (University of Dusseldorf, Germany) based on our previous observations with outcome variations and effect sizes in the mouse model (1, 7, 40, 45, 60). Our sample sizes were determined with the condition of an 80% chance of detecting a moderate effect size. GraphPad Prism V9 (Sorrento Valley, CA) was used for the statistical analysis of data. The data were tested for normality and equality of variance and analyzed using unpaired t-tests given that our data were collected from independent groups. Interactions between samples/groups will be achieved by ANOVA followed by Bonferroni post-hoc tests. Data expressed as mean ± SD, and p < 0.05 considered for statistical significance. Western blot data were quantified by densitometry analysis using ImageJ software (70, 71, 81) normalized to β-actin. The immunostaining intensity or the number of positive cells was quantified using the standard method in ImageJ software (70, 71, 81). We used a double-blinded study design whereby mice were assigned a unique subject number and then randomized to treatment groups in a predetermined manner by a blinded study coordinator. Blinded investigators performed all data acquisition of outcome measures. Following the final data acquisition, mice were decoded, and final analyses were performed.

### Study approval

All animal procedures were conducted in accordance with institutional ethical guidelines for the care and use of laboratory animals, the National Institutes of Health, and the Florida International University Institutional Animal Care and Use Committee (IACUC). All experiments involving CRISPR/Cas9 and AAV-based approaches were conducted in accordance with institutional biosafety requirements and were approved by the Florida International University Institutional Biosafety Committee (IBC).

## Data availability

Data underlying the display items in the manuscript, related to Figures 1, 2, 3, 4, 5, 6, and 7; and S1–S9 are available as Source Data. Microscopy data reported in this paper will be shared by the lead contact upon request. Any additional information required to reanalyze the data reported in this paper is available from the lead contact upon request. All unique reagents generated in this study will be made available from the lead contact (P.M.A.M) and may require a completed materials transfer agreement.

## Supplementary Information

The online version contains supplementary material available at…..

## Author contributions

PMAM: conception; PMAM, SB: experimental design; SB, YAP, SA, PMAM: experimentation and data acquisition; SB, PMAM: data analysis and interpretation; PMAM: figure preparation; SB, PMAM: manuscript writing; PMAM: funding. All authors read and approved the final manuscript.

## Acknowledgements

We thank Anitha Malat and Danielle Caruso at the Core Imaging Facility, Hackensack Meridian Health JFK University Medical Center, Edison, NJ, for their technical assistance. All content was subsequently reviewed and edited by the authors, who take full responsibility for the final version of the manuscript.

