## Supplementary Materials for "Nrf2 regulates ICAM-1–mediated neutrophil extracellular trap formation after traumatic brain injury"

**Supplemental Materials:**

**Supplementary Table 1. Details of the antibodies used for this study.**

| **Antibodies** | **Dilution** | **Catalog Number** | **RRID** | **Vendor** |
| --- | --- | --- | --- | --- |
| Anti-β-actin | WB: 1:1000 | MA575739 | AB_2545348 | ThermoFisher |
| anti-Nrf2 | WB: 1:1000;  IHC: 1:250 | MAB3925 | AB_2263162 | R and D |
| anti-p-Nrf2 | WB: 1:1000;  IHC: 1:250 | PA5-67520 | AB_3353597 | Novus Biologicals |
| anti-GPx1 | WB: 1:1000 | PA5-30593 | AB_2548067 | ThermoFisher |
| anti-GSTm1 | WB: 1:1000 | PA5-22278 | AB_11154815 | ThermoFisher |
| anti-HO-1 | WB: 1:1000 | GTX101147 | AB_1950502 | Gene Tex |
| anti-NQO1 | WB: 1:1000 | ab80588 | AB_1603750 | Abcam |
| anti-NOX1 | WB: 1:1000 | SAB4200097 | AB_10620170 | Sigma- Aldrich |
| anti-4HNE | WB: 1:1000 | ab46545 | AB_722490 | Abcam |
| anti-iNOS | WB: 1:1000 | ab3523 | AB_303872 | Abcam |
| anti-3NT | WB: 1:1000 | 06-284 | AB_310089 | Millipore |
| anti-ICAM-1 | WB: 1:1000;  IHC: 1:250 | MA5407 | AB_223596 | ThermoFisher |
| anti-Mac1 | WB: 1:1000;  IHC: 1:250 | nb11089474 | AB_1216361 | Novus Biologicals |
| anti-LFA 1 | WB: 1:1000;  IHC: 1:250 | ab186873 | Not available | Abcam |
| anti-MMP-2 | WB: 1:1000;  ICC: 1:250 | 87809S | AB_2800107 | Cell Signaling |
| anti-MMP-9 | WB: 1:1000;  ICC: 1:250 | ab76003 | AB_1310463 | Abcam |
| anti-occludin | WB: 1:1000;  ICC/IHC: 1:250 | ab31721 | AB_881773 | Abcam |
| anti-claudin-5 | WB: 1:1000;  ICC/IHC: 1:250 | ab15106 | AB_301652 | Abcam |
| anti-ZO-1 | WB: 1:1000;  ICC/IHC: 1:250 | ab59720 | AB_946249 | Abcam |
| anti-N-cadherin | WB: 1:1000;  ICC: 1:250 | ab18203 | AB_444317 | Abcam |
| anti-connexin-43 | WB: 1:1000 | 3512S | AB_2294590 | Cell Signaling |
| anti-CD68 | IHC: 1:250 | ab955 | AB_307338 | Abcam |
| anti-vWF | ICC/IHC: 1: 250 | Ab11713 | AB_298501 | Abcam |
| anti-Ly6G | WB: 1:1000;  IHC: 1:250 | 87048S | Not available | Cell Signaling |
| anti-H3Cit | WB: 1:1000;  IHC: 1:250 | 17939 | AB_3665810 | Cayman Chemicals |
| anti-H3 | WB: 1:1000 | 9715S | Not available | Cell Signaling |
| anti-PAD4 | WB: 1:1000,  IHC: 1:250 | PA5-22317 | AB_11155990 | ThermoFisher |

ICC: Immunocytochemistry; IHC: immunohistochemistry; WB: western blotting.

**Supplementary Table 2: Primers designed for RT-qPCR expression assays and ChIP-qPCR.**

| **q-PCR primers:** | | |
| --- | --- | --- |
| Gene target | FWD primer sequence (5’🡪 3’) | REV primer sequence (5’🡪 3’) |
| Nrf2 | GCCTTACTCTCCCAGTGAATAC | CTCCCAAATGGTGCCTAAGA |
| GPx1 | GATCTCAGCACCATCCAGTT | GGACAGCAGGGTTTCTATGT |
| GSTm1 | CGCTACATCGCAACACCTAT | GGGTAATTCTAGGAAGCGTGAG |
| HO-1 | CTCCCTGTGTTTCCTTTCTCTC | CAGTCGTGGTCAGTCAACAT |
| NQO1 | AGTGCTCGTAGCAGGATTTG | TCTGGTTGTCAGCTGGAATG |
| GAPDH | GGTCGGTGTGAACGGATTT | GTGGATGCAGGGATGATGTT |
| **ChIP-qPCR primers:** | | |
| Gene target | FWD primer sequence (5’🡪 3’) | REV primer sequence (5’🡪 3’) |
| GPx1 | ACA ATA TAA GGG AGC TGT GCG T | CTA GGG CGG GTC TGG TCT A |
| GSTm1 | GGA CAA AGA AAA GGT GGT ACG | TGG GTT AAC TCA CCC AGA ATG |
| HO-1 | TGA AGT TAA AGC CGT TCC GG | AGC GGC TGG AAT GCT GAG T |
| NQO1 | TCT AAG AGC AGA ACG CAG CA | TTC GTG GGA CCT GCC TAC AT |

**Supplemental Figures:**

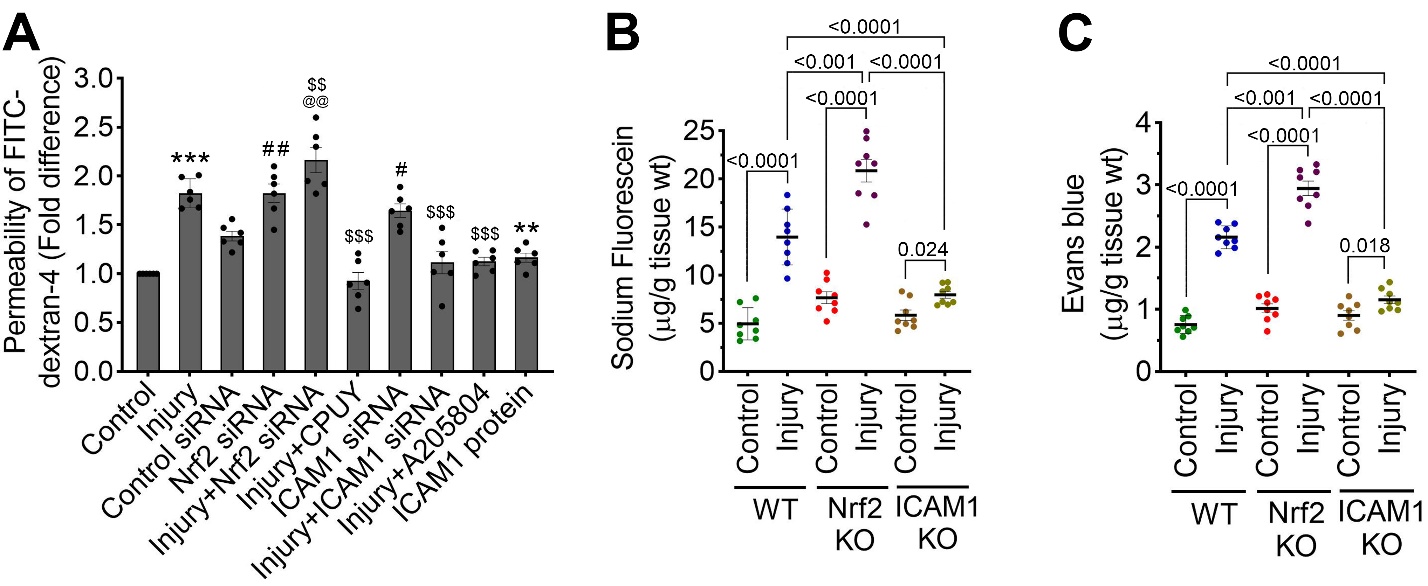

**Supplemental Figure 1. Nrf2 regulates BBB integrity through ICAM-1 following traumatic injury.**

(**A**) *In vitro* assessment of blood–brain barrier (BBB) integrity using FITC-dextran permeability assay in stretch-injured hBMVEC cultures 24 h after treatment with control siRNA, Nrf2 siRNA, the Nrf2 activator CPUY, ICAM-1 siRNA, the ICAM-1 inhibitor A2015804, or recombinant ICAM-1 protein. Bar graphs represent fold change quantification of FITC-dextran-4 permeability normalized to control group (n = 6/group).

(**B-C**) *In vivo* evaluation of BBB permeability following injury using sodium fluorescein (B) and Evans blue (C) tracer assays in wild-type (WT), *Nrf2⁻/⁻,* and *ICAM-1⁻/⁻* mice (n = 8/group).

All values are expressed as mean ± SEM. Statistical analysis was performed using one-way ANOVA in **A** or two-way ANOVA in **B** and **C** followed by Dunnett’s post hoc test.

*p < 0.05* statistically significant. ****P < 0.001* versus control; *^###^P < 0.001* versus control siRNA; *^$$$^P < 0.001* versus injury in **A**.

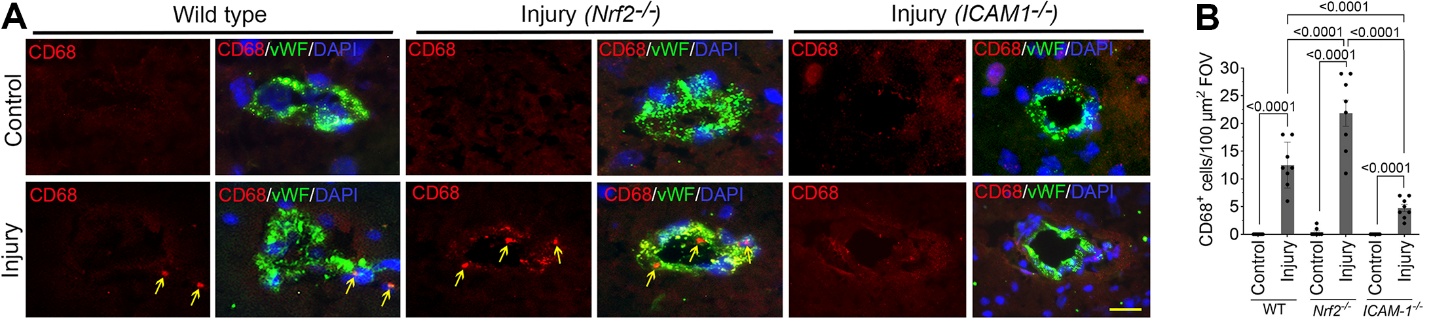

**Supplemental Figure 2. Nrf2 and ICAM-1 regulate macrophage infiltration following in vivo injury.**

(**A-B**) Immunohistochemical analysis of CD68⁺ macrophage (red), and co-localized with von Willibrand Factor (vWF, a microvessel maker, green) and DAPI (nucleus, blue). Infiltration of CD68^+^ cells into the perivascular space in wild-type (WT), *Nrf2⁻/⁻*, and *ICAM-1⁻/⁻* mice under control and injury conditions are shown in the representative images (**A**) and corresponding quantification (**B**) (n = 8/group). Scale bar = 40 µm.

All values are expressed as mean ± SEM. Statistical analysis was performed using two-way ANOVA followed by Dunnett’s post hoc test. *p < 0.05* statistically significant.

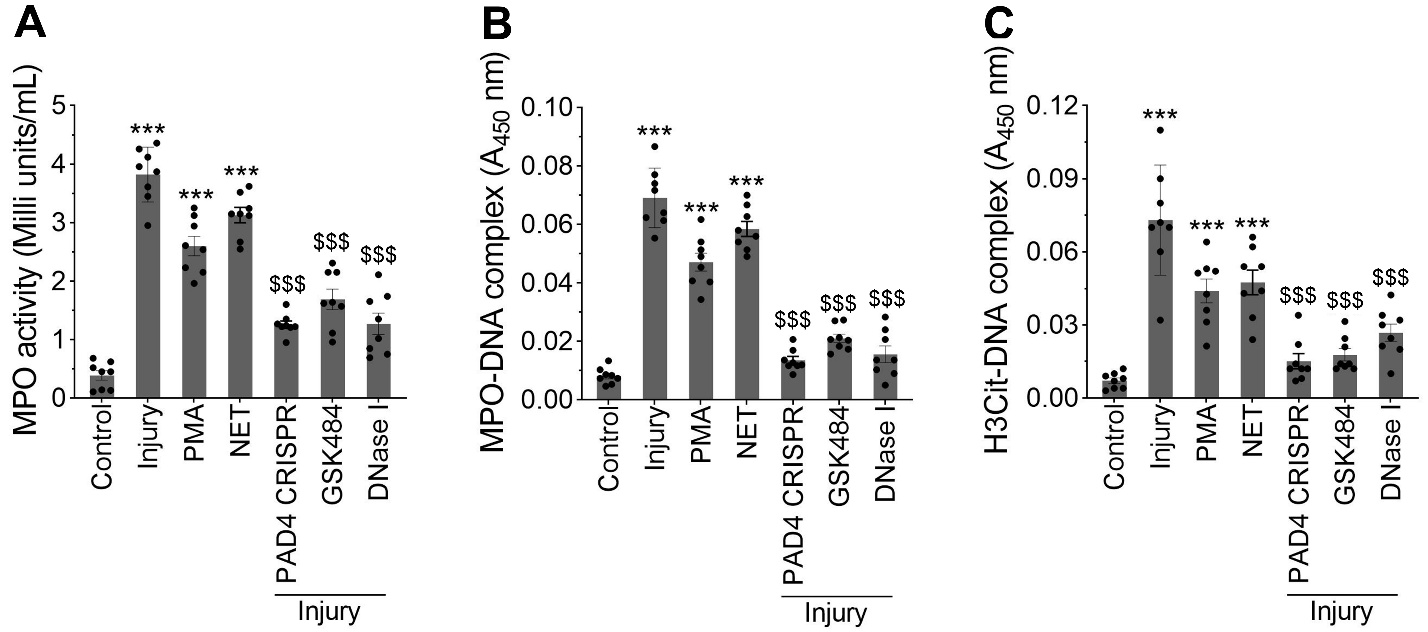

**Supplemental Figure 3. Activation of NET-associated effector pathways in hBMVECs following injury.**

(**A**) Measurement of myeloperoxidase (MPO) activity in human brain microvascular endothelial cells (hBMVECs) under control conditions, injury, and indicated treatments, including PMA stimulation, purified NET exposure, PAD4 CRISPR-mediated inhibition, GSK484 treatment, and DNase I treatment (n = 8/group).

(**B**) Quantification of MPO-DNA complex formation as an indicator of neutrophil extracellular trap (NET) activity across experimental groups (n = 8/group).

(**C**) Quantification of citrullinated histone H3 (H3Cit)-DNA complexes in the same conditions (n = 8/group).

Data are presented as individual data points with mean ± SEM. Statistical analysis was performed using one-way ANOVA followed by Dunnett’s post hoc test. *p < 0.05* statistically significant. ****P < 0.001* versus control; *^$$$^P < 0.001* versus injury.

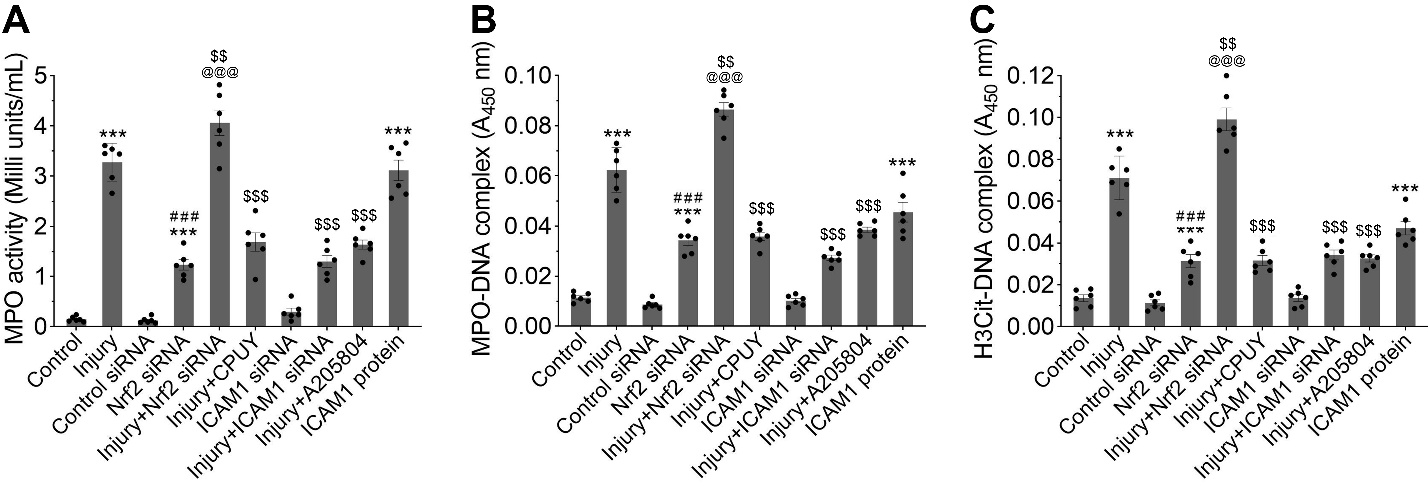

**Supplemental Figure 4. Nrf2 and ICAM-1 regulate NET formation following injury in hBMVECs.**

(**A**) Measurement of myeloperoxidase (MPO) activity in human brain microvascular endothelial cells (hBMVECs) under control and injury conditions, with Nrf2 knockdown (siRNA), Nrf2 activation (CPUY192018), ICAM-1 knockdown (siRNA), ICAM-1 inhibition (A205804), and recombinant ICAM-1 protein treatment (n = 6/group).

(**B**) Quantification of MPO-DNA complex formation across the indicated experimental conditions (n = 6/group).

(**C**) Quantification of citrullinated histone H3 (H3Cit)-DNA complexes as a marker of neutrophil extracellular trap (NET) formation (n = 6/group).

All values are expressed as mean ± SEM. Statistical analysis was performed using one-way ANOVA followed by Dunnett’s post hoc test and statistical significance between groups is indicated on the graphs. *p < 0.05* statistically significant. ****P < 0.001* versus control; *^###^P < 0.001* versus control siRNA; *^$$$^P < 0.001* versus injury.

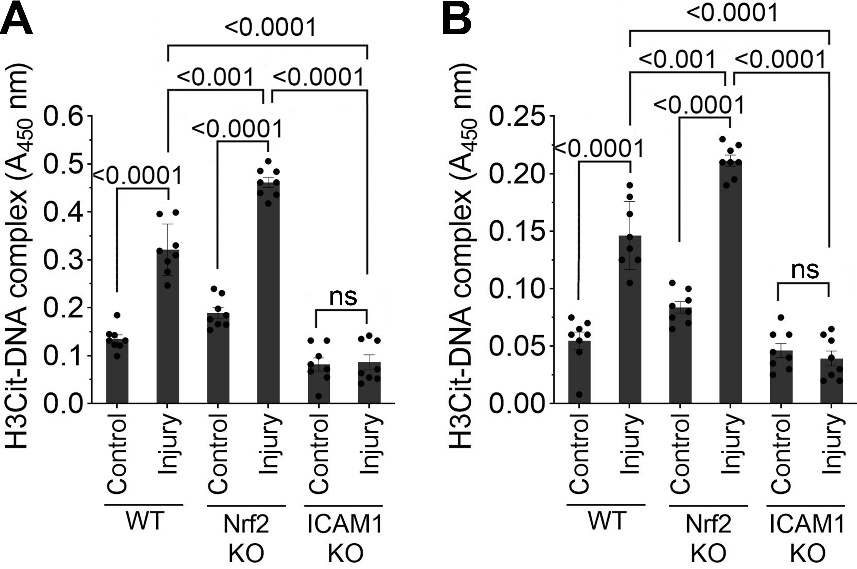

**Supplemental Figure 5. Nrf2 and ICAM-1 regulate circulating and tissue NET formation following *in vivo* injury.**

(**A-B**) Quantification of citrullinated histone H3 (H3Cit)-DNA complexes by ELISA in brain tissue lysates (**A**) and plasma samples (**B**) from wild-type (WT), *Nrf2⁻/⁻*, and *ICAM-1⁻/⁻* mice under control and injury conditions (n = 8/group).

All values are expressed as mean ± SEM. Statistical analysis was performed using two-way ANOVA followed by Dunnett’s post hoc test. *p < 0.05* statistically significant.

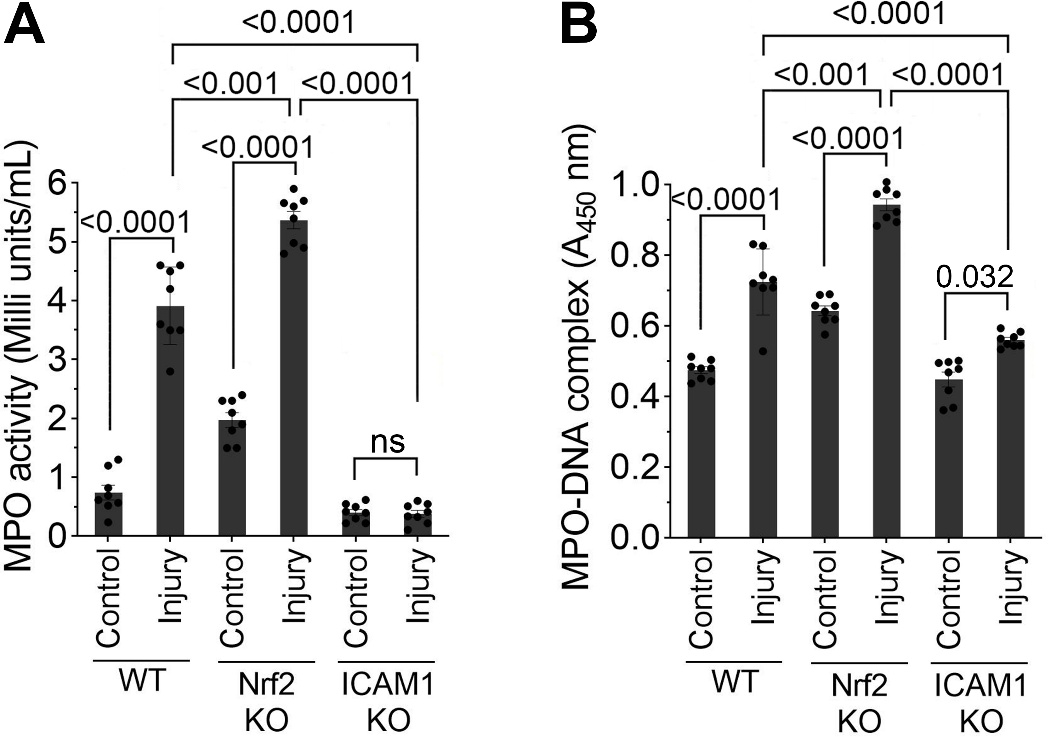

**Supplemental Figure 6. Nrf2 and ICAM-1 regulate MPO activity and NET-associated MPO release following *in vivo* injury.**

(**A**) Measurement of myeloperoxidase (MPO) activity in brain tissue lysates from wild-type (WT), *Nrf2⁻/⁻*, and *ICAM-1⁻/⁻* mice under control and injury conditions (n = 8/group).

(**B**) Quantification of MPO-DNA complexes in plasma samples as an indicator of NET-associated MPO release across the same experimental groups (n = 8/group).

All values are expressed as mean ± SEM. Statistical analysis was performed using two-way ANOVA followed by Dunnett’s post hoc test. *p < 0.05* statistically significant.
